# Population dynamics after pancreatitis dictates long-lasting epigenetic reprogramming and mediates tumor predisposition

**DOI:** 10.1101/2024.07.04.600210

**Authors:** Marco Fantuz, Johannes Liebig, Giovanni Fontana, Valerio Iebba, Carmine Carbone, Sören Lukassen, Christian Conrad, Alessandro Carrer

## Abstract

Local inflammation in the pancreas is transient but imprints a durable epigenetic memory on epithelial cells, making them more amenable to oncogenic transformation. However, it is unclear whether epithelial cell heterogeneity is impacted by acute pancreatitis (AP) or whether population dynamics during regeneration contributes to the establishment of inflammation memory.

To tackle those questions, we deployed experimental pancreatitis in mice and performed paired sequencing of transcriptomic and chromatin accessibility profiles at single nucleus resolution. We documented cell type abundance but also applied integrative analyses to infer phenotypically-distinct clusters of mesenchymal and exocrine cells. We found that AP perturbs a subset of “idling” acinar cells, which separate from more canonical “secretory” acini based on a more diversified proteome, which include elevated expression of signal transduction receptors. We linked acinar cell heterogeneity to epigenetic differences that also endow idling cells with superior plasticity. These constitute about 40% of acinar cells but can proliferate and skew their phenotype in response to AP. This leads to a remarkable recovery of pancreas histology and function, but also to the dissemination of idling-like features across the exocrine parenchyma. Mechanistically, idling acinar cells are characterized by enhanced transcriptional activity and protein synthesis. After recovery from pancreatitis, acini show elevation of both and establishment of chronic Unfolded Protein Response (UPR). We finally demonstrated that AP-primed pancreata show signs of elevated UPR and that ER stress promotes acinar cell metaplasia.

Our data interrogate phenotypical dynamics during tissue regeneration to identify cell states amenable to epigenetic imprinting. They also suggest that UPR-alleviating strategies might curtail the risk of developing pancreatic cancer for individuals who experiences AP.

## Introduction

Pioneering work dating back in the 50s and 60s established the ability of pancreatic tissue to regenerate and quickly restore normal function and histology after injury^1^. However, pancreata previously exposed to acute pancreatitis (AP) develop cancer more quickly upon induction of oncogenic *Kras*^2,3^. Retrospective analyses showed that individuals who suffered AP are at elevated risk of pancreatic cancer, even several decades after the episode^4,5^. These observations strongly suggest that AP elicits an incomplete regenerative process where epithelial cells retain memory of past inflammatory episodes. Understanding cellular and molecular dynamics of pancreatic regeneration is critical to avoid permanent disturbances that might elevate cancer risk.

The pancreas is organized in exocrine and endocrine compartments, the former being markedly more abundant and able to regenerate more efficiently^6^. Acinar cells, which exert synthetic and secretory functions, and ductal cells, which line to form columnar epithelia and tree-branching conducts for the draining of pancreatic juice, together constitute the exocrine compartment. They share common developmental progenitors and acinar cells can transdifferentiate into ductal cells, a process associated to tissue repair, termed acinar-to-ductal metaplasia (ADM)^10,11^. Single cell- and nucleus sequencing unveiled significant heterogeneity of both cell types^12–15^, although it is not clear how subpopulation balance is impacted during tissue regeneration or whether specific subtypes are endowed with superior plasticity.

Pancreatitis is a painful inflammation of the pancreas that represents a common cause of hospitalization. When acute, pancreatitis is transient in that pancreatic functions are spontaneously regained when aetiological insults are alleviated^16^. Around 65–70% of patients with acute pancreatitis (AP) are discharged without complications, with resolution of symptoms within few days^17^. Multiple experimental models in rodents have been developed; they show that AP causes abundant loss of acinar cells, initiation of ADM and remodeling of stromal compartment but also demonstrate complete histological restoration^2,18^, which also likely to occur in humans. Among those models, the most commonly adopted is supraphysiological stimulation of acinar cell secretion by cerulein, an analogue of the hormone cholecystokinin^19,20^. Following cerulein injection, acinar cells dynamically reprogram their epigenome^21^ to modulate the binding of pioneer transcription factors and the activity of lineage-dependent regulatory proteins^22–24^. Murine models also demonstrated that pancreatitis cooperates with oncogenic KRAS to induce epigenetic states in epithelial cells that unleash cell plasticity and initiate carcinogenesis^25–27^. Together, data suggest that AP elicits actionable acinar cell states that facilitate regeneration, but also make cells vulnerable to oncogenic transformation.

It was also demonstrated that the epigenomic landscape of acinar cells can be permanently remodeled by AP, with increased markers of active chromatin at “memory” loci and lowered epigenetic barriers to oncogenic transformation even several weeks after AP induction^2,3^. This aligns with the notion that epithelial cells can be primed by inflammation and can retain pro-inflammatory or pro-oncogenic features^28,29^. Yet, it remains unclear how the epigenetic memory is established at specific loci. In addition, mechanisms linking a perturbed epigenome to facilitated carcinogenesis are incompletely understood.

We speculated that phenotypic plasticity of acinar cells contributes to the establishment of an inflammation memory and to cancer predisposition. To tackle this issue, we performed paired single nucleus RNA and ATAC sequencing (sn-RNA/ATAC-seq) to document enduring phenotypic transitions induced by experimental AP in mice. Our approach allowed to reconstruct subpopulation dynamics and identify a subset of “idling” acinar cells endowed with superior plasticity. This aligns with - and complement - our single nucleus topography of the human pancreas^13^. We also documented widespread chromatin opening that initiates at the “idling” subset of acinar cells and eventually propagates to all acini, inducing chronic proteostatic stress. Combining bioinformatic modeling and classical biochemistry we found that protein dyshomeostasis facilitates ADM and promotes tumor formation. Our data indicate that memory of AP resides at the level of acinar cells and evolves after resolution of inflammation as consequence – at least in part – of acinar subpopulation dynamics.

## Results

We deployed a widely described experimental set-up to induce transient inflammation of murine pancreas through iterative injections of cerulein over two consecutive days^30^. Wounding occurs when cerulein administration stops, and injury is fully repaired without scars^19^. Transmission electron microscopy (TEM) showed that the structure of acini – the glomerular clusters of cells that constitute the functional unit of exocrine pancreas – is indeed not impacted by prior inflammation (Figure 1B). In line, exocrine function was not perturbed one month after cerulein injection (Suppl. Figure 1A). Also, no signs of cell replication could be observed at this time point (Suppl. Figure 1B).

**Figure 1:**
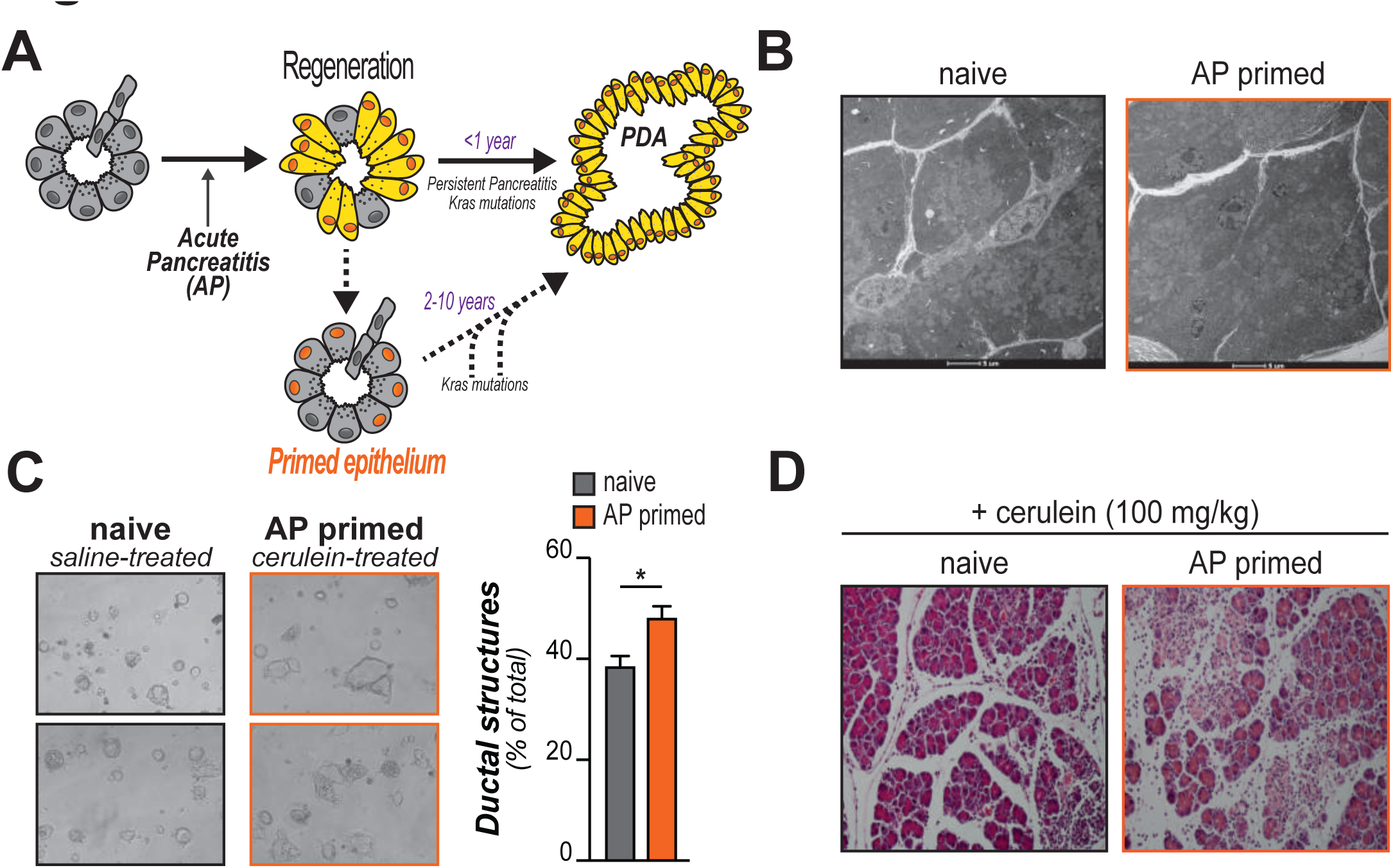
histological normal pancreata can be primed by acute pancreatitis (AP). **A**, working hypothesis, schematic. Acute pancreatitis (AP) induces an inflammatory reaction and tissue regeneration, which is associated to phenotypical and morphological plasticity of the epithelium. Oncogenes co-opt these states to initiate tumorigenesis. Indeed, pancreatic ductal adenocarcinoma (PDA) arises soon after AP episodes in humans (highlighted in purple). In most cases however, tissue damage is resolved with no obvious consequences. Data suggest that the pancreatic epithelium is however permanently primed by AP episodes and remain poised for tumorigenesis even after several years. **B-D,** mice were injected with cerulein for 2 consecutive days and were then allowed to recover for additional 28 days. Control mice were injected in parallel with saline. **B** shows transient pancreatitis transmission electron microscopy (TEM) of acini structure (representative images; *n=3 mice/group*). In **C**, acinar explants were purified and immediately embedded in Matrigel®. After two days, cytological images were acquired and examined for the presence of duct-like structures (*n=3* mice/group; *>30* images each prep). In **D**, mice were subsequently injected with cerulein (8 hourly injections) and sacrificed the morning after. The figure shows representative sections stained with hematoxylin and eosin.

Isolated acinar explants undergo spontaneous metaplasia when cultured in Matrigel and the number/size of duct-like structures correlates with cell plasticity and susceptibility to oncogenic transformation^10^. Interestingly, we observed that acute pancreatitis (AP)-primed acinar cells exhibited enhanced plasticity (Figure 1C). In addition, mice that had undergone inflammation/regeneration showed elevated sensitivity to cerulein re-administration that elicits a more abundant infiltration of mononuclear cells in AP-primed pancreata (Figure 1D).

We asked whether this might be due to incomplete or impaired homeostatic regeneration. Histological examination showed widespread inflammation acutely after cerulein administration (1-3 days), with extensive immune infiltration, dilation of interstitial space and regeneration-associated metaplasia. However, tissue architecture was restored after one week (hereafter termed, *post-AP early*) and preserved at longer time points (hereafter termed, *post-AP late* - four weeks after cerulein injection) (time point histological examination in: Suppl. Figure 1C). No remarkable differences in tissue architecture (number of blood vessels or duct, assessed by staining with CD31 or KRT19 respectively) or immune cell infiltration were observed *early* or *late* post AP (Suppl. Figure 1D).

These findings indicate that AP episodes endow acinar cells with superior plasticity and sensitize to late insults. These changes are long lasting but are not associated with overt changes in pancreatic cytology, histology or function.

### Single nucleus profiling of tissue homeostasis after AP

We reasoned that AP primes acinar cells to increase cancer susceptibility through multiple, possibly co-existing, mechanisms that enable a more permissive environment even after inflammation is resolved. These include functional remodeling of the stromal population (“microenvironmental memory”), enrichment in progenitor-like subpopulations (“cellular memory”) and permanent reprogramming of the epigenome (“molecular memory”). These have all been shown to play a role in PDA carcinogenesis and are summarized in Figure 2A.

**Figure 2:**
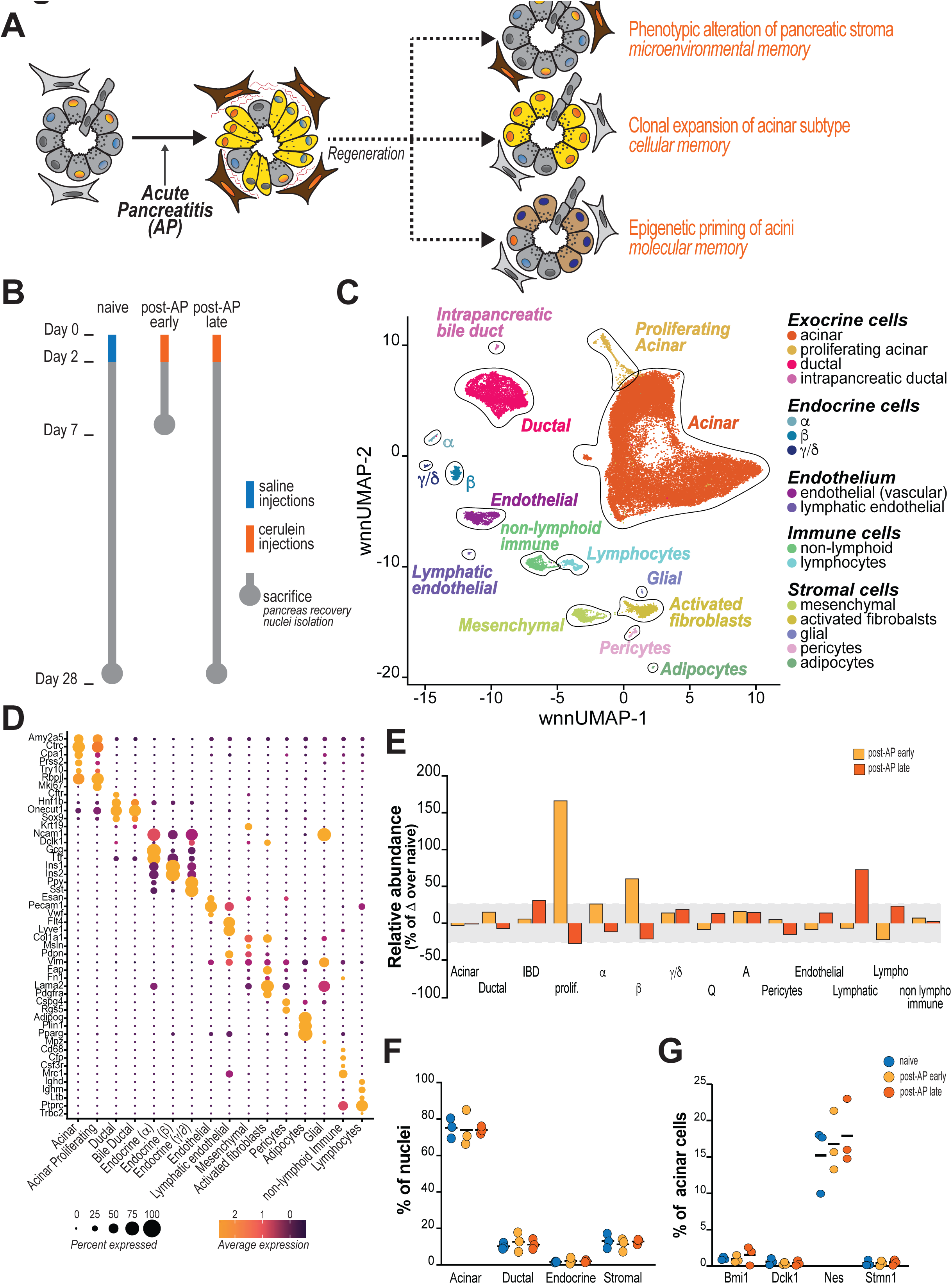
paired single nucleus analysis for the characterization of pancreatitis memory. **A**, hypotheses for mechanisms of tissue memory after inflammation. **B,** schematic of experimental design and samples analyzed. **C,** Weighted shared nearest neighborhood of uniform manifold approximation and projection (wnnUMAP) of paired RNA- and ATAC-sequencing of nuclei isolated from mice as in B (*n=3* mice/condition). Clusters are defined by solid black lines and highlighted with different colors. Color legend on the right. **D,** Dot-plot showing the expression of lineage-associated marker genes. **E,** abundance of identified clusters was quantified in naïve, post-AP *early* (7 days) and post-AP *late* (28 days) samples. For post-AP samples, difference with naïve (expressed as %) was plotted for each population (adipocytes were removed due to very low abundance). **F,** Abundance of exocrine (acinar and ductal cells, split), endocrine and stromal cells was plotted. Each dot denotes a mouse. **G,** Abundance of known progenitor cells was plotted. Each dot denotes a mouse.

To interrogate and parse these plausible mechanisms, we performed granular analysis of both population dynamics and epigenomic reprogramming applying single nucleus RNA/ATAC-Seq that simultaneously map gene expression and chromatin accessibility in isolated nuclei. Mice were injected with either saline or cerulein and pancreata snap frozen at either 7 or 28 days post AP induction (n=3 mice each condition), when tissue regeneration is completed and tissue architecture fully restored (Figure 2B, Suppl. Figure 1C). A total of 32329 nuclei (median: 3145 per mouse; median interquartile range 2822-4598 cells/mouse) were purified applying a recently-optimized protocol^13^ and RNA/ATAC libraries sequenced. We applied a weighted analysis of shared nearest neighborhoods (WNN) to balance differences in sequencing depth and peak distribution^31^ and to finally identify 22 clusters that were associated to 15 different cell types based on the expression of well-characterized identity markers (Figure 2C-D). While gene expression provided a more granular separation of cell populations, chromatin accessibility consolidated lineage-related cell types in fewer and larger clusters (i.e.: acinar cells, immune cells, blood/lymph endothelial cells) (Suppl. Figure 2A). This observation aligns with the notion that chromatin elements dictates cell identity, including for pancreatic cells^32^. Of note, our approach allows the efficient recovery of exocrine cells, which constitute the largest cell population; this is in striking contrast with previously documented single cell sequencing datasets where the exocrine compartment is markedly underrepresented due to rapid RNA degradation (Suppl. Figure 2B). In addition, our approach avoids common procedural steps (cell dissociation, FACS sorting) that significantly perturb the epigenome^33–35^. For context, exocrine cell gene expression peaks largely overlapped with prior analyses of sorted epithelial cells, while patterns of chromatin accessibility were notably different (Suppl. Figure 2C). We ascribe those differences to the fact that we could work from rapidly-frozen and intact nuclei. Yet, peaks at major lineage identity-related loci were consistently found and highly reproducible across independent biological replicates (Suppl. Figure 2D). We conclude that our analysis efficiently examines unperturbed nuclei and inherently unites multiple sequencing approaches, avoiding the introduction of procedural biases and bypassing the need for *in silico* superimposition of independently-sequenced samples.

We proceeded to seek whether AP induces persistent perturbations of tissue composition. In fact, some less-represented clusters were affected by AP albeit variations were negligible at post-AP late time point (Figure 2E). Abundance of exocrine (both acinar and ductal), endocrine and stromal cells is not impacted by prior inflammation and cell count is homeostatically restored already at 7 days post AP (“post-AP early”) (Figure 2F). Of note, we could not detect Tuft cells (*Pouf23+*/*Dclk1+*) in our dataset (in accord with: ^13,36^), although we observed the presence of intrapancreatic biliary ductal cells (*Cftr*-/*Hnf1b*+). This aligns with prior analysis of ductal cell heterogeneity in normal pancreas^12^. We also did not found evidence for metaplastic cells, documented in single cell analyses of either chronic pancreatitis or mutant KRAS-induced metaplasia^36–38^. Similarly, abundance of progenitor-like populations (*Bmi*+, *Dclk1*+ and *Stmn1*+; REF:^8^) was limited and not significantly altered by AP in our analysis (Figure 2G; Suppl. Figure 2E). *Nes*+ cells were more abundant but again their number or distribution was not affected by AP (Figure 2G; Suppl. Figure 2E). This confirms full resolution of local inflammation and complete restoration of tissue organization without overt misrepresentation of distinct progenitor cell populations. Importantly, in-depth analysis was performed on unfiltered DNA fragments and we detected no evidence for non-mammalian genomes (pancreas-infiltrating bacteria or yeasts) nor AP-induced mutations at pro-oncogenic genes, including *Kras*, *Tp53*, *Smad4*, *Cdkn2a*^39^ (*data not shown*). These results corroborate full restoration of normal pancreatic histology observed in Figure 1.

However, proliferating acinar cells (positive for *Mki67* expression) were markedly more abundant in *post-AP early* (but not *late*) mice; this was confirmed by immunostaining of formalin-fixed tissues (Suppl. Figure 2F-G). Only a small fraction of these cells express known markers of pancreatic progenitors (*Nes* and/or *Stmn1*) or genes denoting ADM (Suppl. Figure 2H). Compared to non-duplicating acinar cells, their expression profile is enriched for genes required for pancreatic regeneration (Suppl. Figure 2I).

We conclude that our analysis profiles the cellular composition of mouse pancreata at high resolution and identifies all cell populations in the pancreatic milieu, including a cluster of proliferating acinar cells. These are *bona fide* mature acini undergoing fission and likely contribute to tissue homeostasis post AP, as previously demonstrated using a continuous cell-tracing strategy^9^. Their elevated abundance at earlier time points indicates incomplete regeneration despite apparent histological normalcy, but also suggests that they unlikely contribute to enduring tumor susceptibility as the number of proliferating acinar cells eventually returns to normal.

### Resident mesenchymal cells remain activated after resolution of pancreatitis

Stromal cells are highly plastic and can assume multiple phenotypic states that can exert heterogeneous cell-extrinsic effects in the tumor/tissue microenvironment. Because inflammation can skew their specification, we looked at fibroblast-like cells, which are abundant in our dataset. We found that the collagen-producing mesenchymal population (*Msln*+, *Vim*+, *Pdpn*+, *Itgb1*+) can be divided in two subgroups: quiescent (Q) and activated (A) (Figure 2C-D). The former is likely constituted by both pancreatic stellate cells and resident fibroblasts while the latter include activated fibroblasts and shows a clear trend to augment post-AP (Figure 3A), suggesting the possibility that these might maintain memory of inflammation (*mechanism #1 in Figure 2A*).

**Figure 3:**
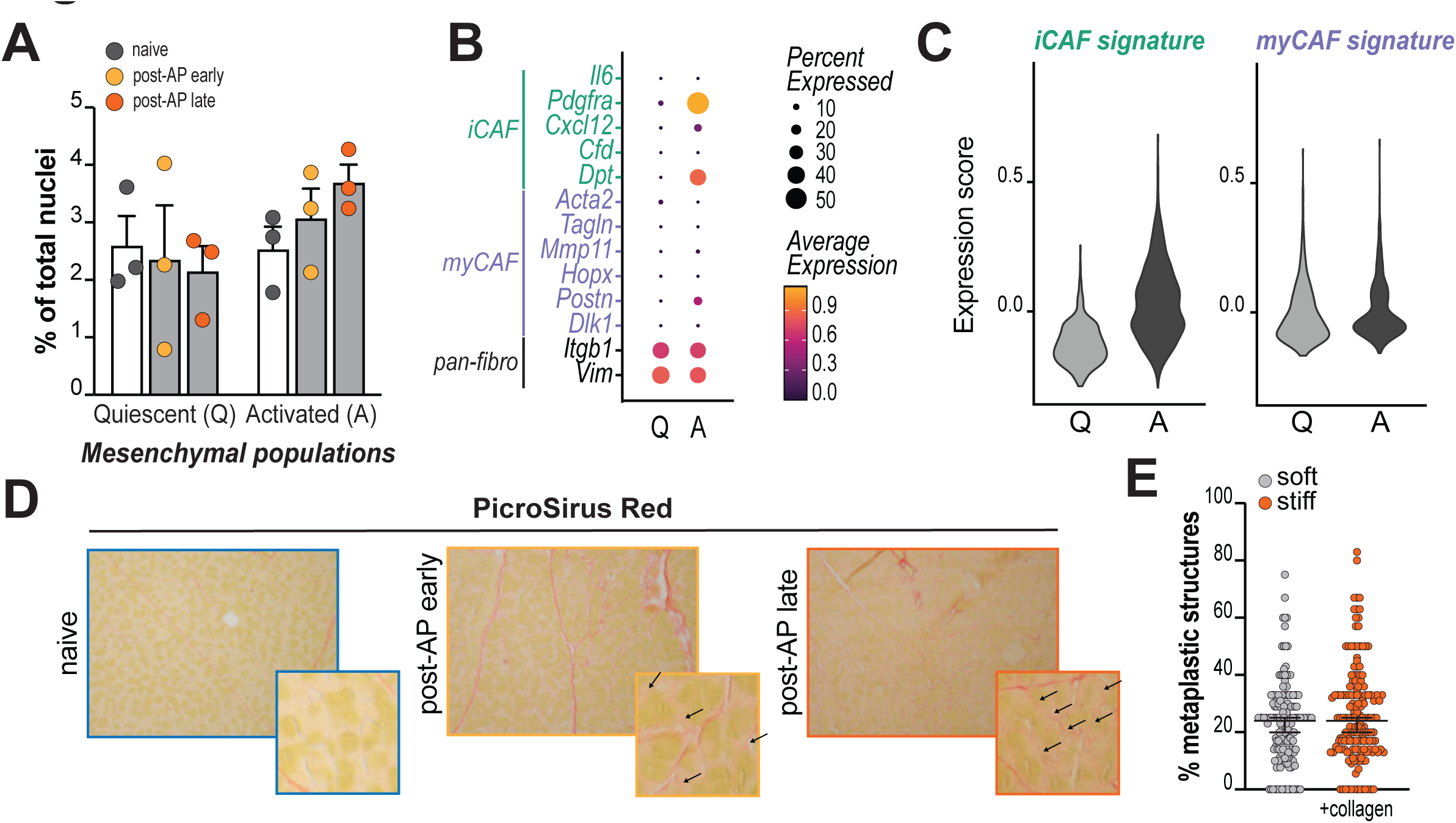
reprogramming of pancreatic mesenchyme after AP. **A**, abundance of non-activated mesenchymal cells (quiescent; Q) and activated fibroblasts (A) at different time points after AP induction. Each dot denotes a mouse. Values are expressed as percentage of total nuclei analyzed. **B,** Dot-plot showing the expression of genes universally expressed in tissue-resident fibroblasts (pan-fibro) or in alternatively-activated subpopulation of cancer-associated fibroblasts (myofibroblast-like – myCAF – or inflammatory – iCAF). **C,** Violin plot showing the expression score of myCAF or iCAF gene signatures (*see Methods*) in quiescent (Q) or activated (A) mesenchymal cells. **D,** PicroSirus Red staining of pancreatic tissue sections from naïve or AP-primed mice (see Figure 2B for timepoints). Representative images of *n=3* mice/group. **E,** acinar explants were purified and immediately embedded in either pure Matrigel® or a 1:1 dilution with rat Collagen I. After two days, cytological images were acquired and examined for the presence of duct-like structures (*n=3* mice/group; *>30* images each prep). Dots denote analyzed optical fields.

Notwithstanding their heterogeneous nature, activated fibroblasts in our dataset are negative for α-SMA and collectively show a quasi-inflammatory phenotype (iCAF-like) (Figure 3B-C; Suppl. Figure 3A) that has been associated to tumor promotion^40^. Notably, pre- and post-AP activated fibroblasts have identical epigenetic profiles (no differentially-accessible loci in naïve vs post-AP fibroblasts), indicating they are not subject to immunological training.

In an attempt to define how activated fibroblasts could facilitate pancreatic carcinogenesis, we observed a thickening of the basal lamina around acini in post-AP mice (Figure 3D), indicative of sustained collagen deposition and augmented stiffness of the extracellular matrix. We thus asked whether moderate changes in tissue elasticity might contribute to acinar cell plasticity and eventually cooperate with oncogenes to drive tumor initiation. However, we found that increasing the elastic modulus to a Matrigel-based 3D matrix does not impact acinar-to-ductal metaplasia of primary wild-type acinar explants (Figure 3E). These findings indicate that subtle but permanent alterations to pancreatic architecture establish after inflammation but do not markedly contribute to AP-linked tumor predisposition.

### Acinar cells are phenotypically heterogeneous

We next applied a tree-aggregated Bayesian modeling (tascCODA^41^) specifically designed for differential population analysis in composite systems. This model uses hierarchical ordering to infer compositional shifts of cellular subtypes weighting the impact of each covariate on each population. The analysis provides an unbiased examination of clusters dynamics and highlights statistically robust changes in cellular subtype abundance. We analyzed the 22 clusters identified and pre-set hierarchical ordering based on cellular lineage (exocrine, endocrine, mesenchymal, immune). Results clearly show that the acinar is the most dynamic cell population following pancreatitis with several statistically-significant changes in the relative abundance of its subclusters (Figure 4A).

**Figure 4:**
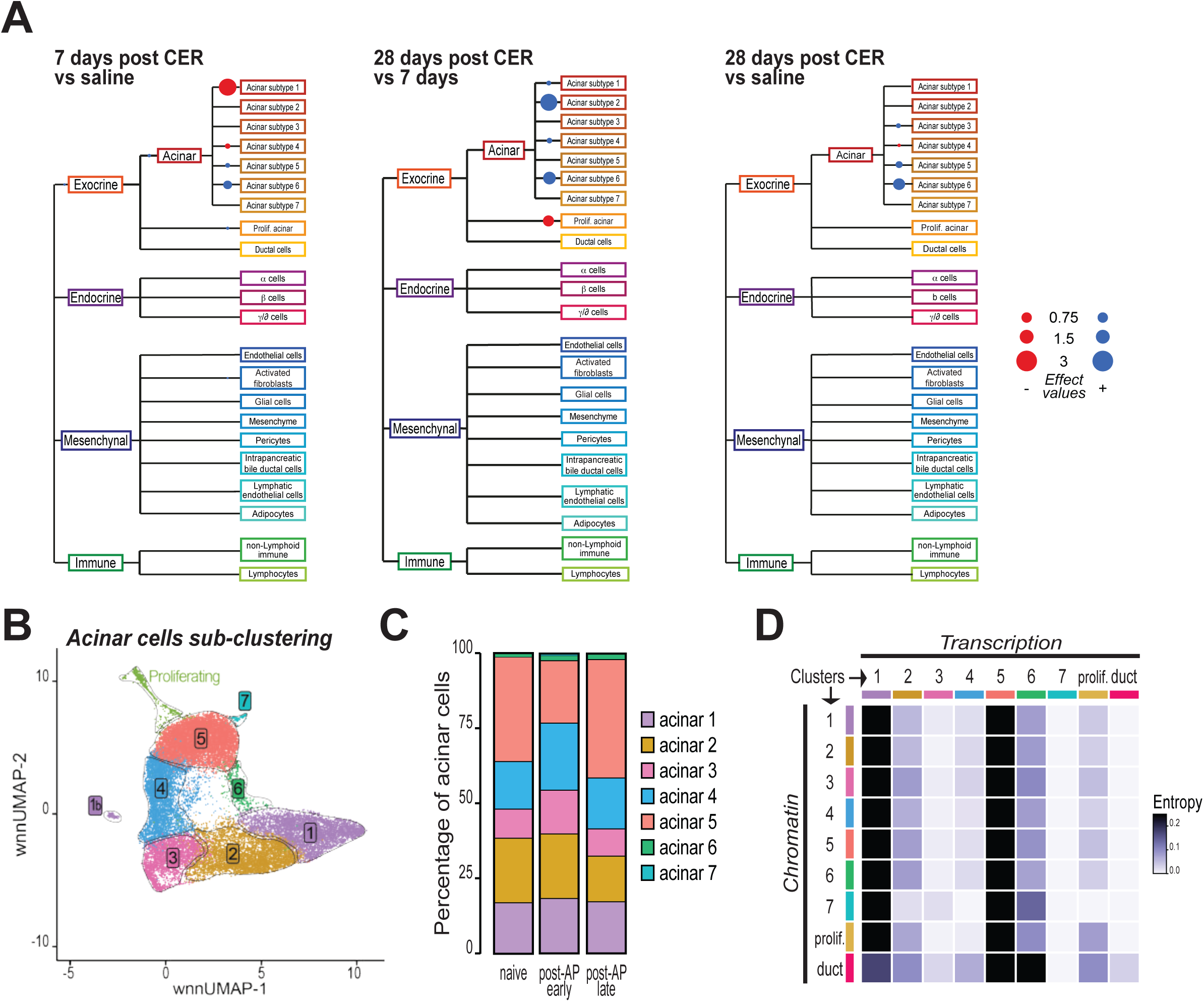
acinar cells exhibit multiple phenotypes. **A**, tascCODA analysis (*see Methods*) of sn-RNA/ATAC-seq clusters, repeated against three possible time-matched combinations. Hierarchy was assigned based on developmental origin and highlighted by color coding. Dot along tree lines denote statistically-significant effect values (legend on the far right). **B,** wnnUMAP plot of acinar cell clustering. Clusters highlighted by different colors and marked by solid black lines. **C,** Acinar subclusters abundance for each time point analyzed. Values are expressed as percentage of acinar nuclei analyzed at for each condition. Color-coded bars show aggregate numbers of 3 independent replicates (mice). **D,** Classifier confusion matrix based on paired sequencing. Values represent number of nuclei from an epigenomic cluster that classify to a transcriptomic cluster, normalized within each row. Black boxes highlight high plasticity epigenomic states.

This indicates that the exocrine compartment – and acinar cells in particular – is sensitive to AP events. For instance, proliferating acinar cells were shown to be specifically enriched at 7 days post cerulein administration (Figure 4A, left and middle panels), confirming our previous observations (Suppl. Figure 2F-G). We could also clearly observe subpopulation remodeling after 7 days (between post-AP *early* and *late*) with enrichment of certain clusters (i.e.: acinar clusters denoted by blue dots). This suggests that, in spite of full histological recovery, tissue homeostasis is incomplete one week after AP induction and regenerative process further remodels the composition of the exocrine parenchyma.

WNN analysis identified 7 clusters of post-mitotic acinar cells, and one additional cluster assigned to proliferating acinar cells (Figure 4B). Relative abundance of acinar clusters changes upon pancreatitis, with marked changes observed “*early*”: contraction of cluster “5”, expansion of clusters “6” and “7” (Suppl. Figure 4A). Yet, cluster composition was largely homeostatic and relative abundances were not significantly altered at “*late*” time point although cluster “6” showed a tendency to remain elevated (Figure 4C, Suppl. Figure 4A). In line, cluster “6” was the most consistently deregulated in tascCODA analysis (Figure 4A – across panels). This indicates that AP perturbs homeostatic equilibrium of pancreatic epithelial cells and dyshomeostasis persists even when normal histology is restored (“post-AP *early*”, 7 days). However, balance of acinar cell subsets is near-completely restored at later time points (“post-AP *late*”, 28 days) leading to the conclusion that clonal expansion of AP-resistant acinar cells does not occur and cannot explain long-lasting memory of inflammation, as previously suggested^15^.

Of note, single cell RNA atlas of human pancreata could characterize only 3 acinar cell subpopulations (secretory, idling and REG+: ^13^), suggesting that chromatin organization pinpoints heightened heterogeneity – at least in mouse. In line, analysis of the correlation among chromatin conformations revealed clear separation of acinar cell subclusters (Suppl. Figure 4B). On the other side, transcriptional profiles of acinar subsets exhibited more vicinity, with clusters (“2”, “3”, “4”) and (“5”, “7”) being closely related (Suppl. Figure 4B).

These findings suggest that acinar cells exhibit phenotypical heterogeneity, generated by the dealignment between chromatin accessibility and gene expression. To better link acinar cell plasticity to relaxed epigenetic boundaries, we applied an algorithm to quantify the degree of entropy between chromatin landscapes and their transcriptional outputs^26,42^. In other words, we measured the phenotypic infidelity of acinar cell subclusters based on the poor coordination between chromatin accessibility at a given locus and transcription of proximal gene(s). This analysis showed that clusters “1” and “5” represent highly plastic states, with potential to cover the full spectrum of acinar cell phenotypes and even transdifferentiate into duct cells (Figure 4D). Interestingly, our model also predicts that acinar cluster “6” exhibits the ability to preferentially evolve into duct cells (Figure 4E). We conclude that the chromatin landscape makes acinar cells phenotypically plastic and anticipate that certain states predispose cells to metaplastic transition.

### Secretory and idling acini are homeostatically regulated in response to pancreatitis

We set out to better characterize the dynamic response of acinar cell states to AP. First, we contextualized our findings with prior work that documented pancreas topography^13^. Human acinar cells exhibit phenotypical dichotomy, and can be classified into two functional states:

“secretory” (*S*) or “idling” (*I*) ^13^. The former are characterized by heightened expression of digestive enzymes, the latter by expression of receptor and signaling mediators that make them reactive to microenvironmental stimuli. Interestingly, cluster “1” in our dataset shows marked overlap with (*S*) acinar cell transcriptional profile but poor similarity with (*I*) transcriptome; cluster “5” shows opposite behavior (Figure 5A). This indicates that secretory and idling subtypes of acinar cells can be detected also in murine pancreata, but also confirms the existence of additional – possibly corollary – states. The RNA expression correlation matrix clearly showed that cluster “5” can be consolidated into a slightly larger group that also contains cluster “7”; interestingly, the two also show a markedly “idling” phenotype (Suppl. Figure 5A) and are separated essentially by differences in chromatin conformation (Suppl. Figure 4B). Similarly, clusters “2”, “3” and “4” are highly related and can be grouped together, while cluster “1” and cluster “6” are clearly distinct (Suppl. Figure 4B – metagroups delimited by orange lines). We termed the congregate cluster [2+3+4] “transitional (*T*) acinar”. Interestingly, the transcriptional profile of cluster “6” overlaps both (*S*) and (*I*) acinar cells and are thus termed “hybrid (*H*) acinar” (Figure 5A). Together, our approach identified four “*metaclusters*” of phenotypically distinct acinar cells: secretory, idling, hybrid and transitional (Suppl. Figure 5A-B). Of note, these acinar populations are clearly perturbed by AP but homeostatically return to a naïve-like equilibrium (Figure 5B).

**Figure 5:**
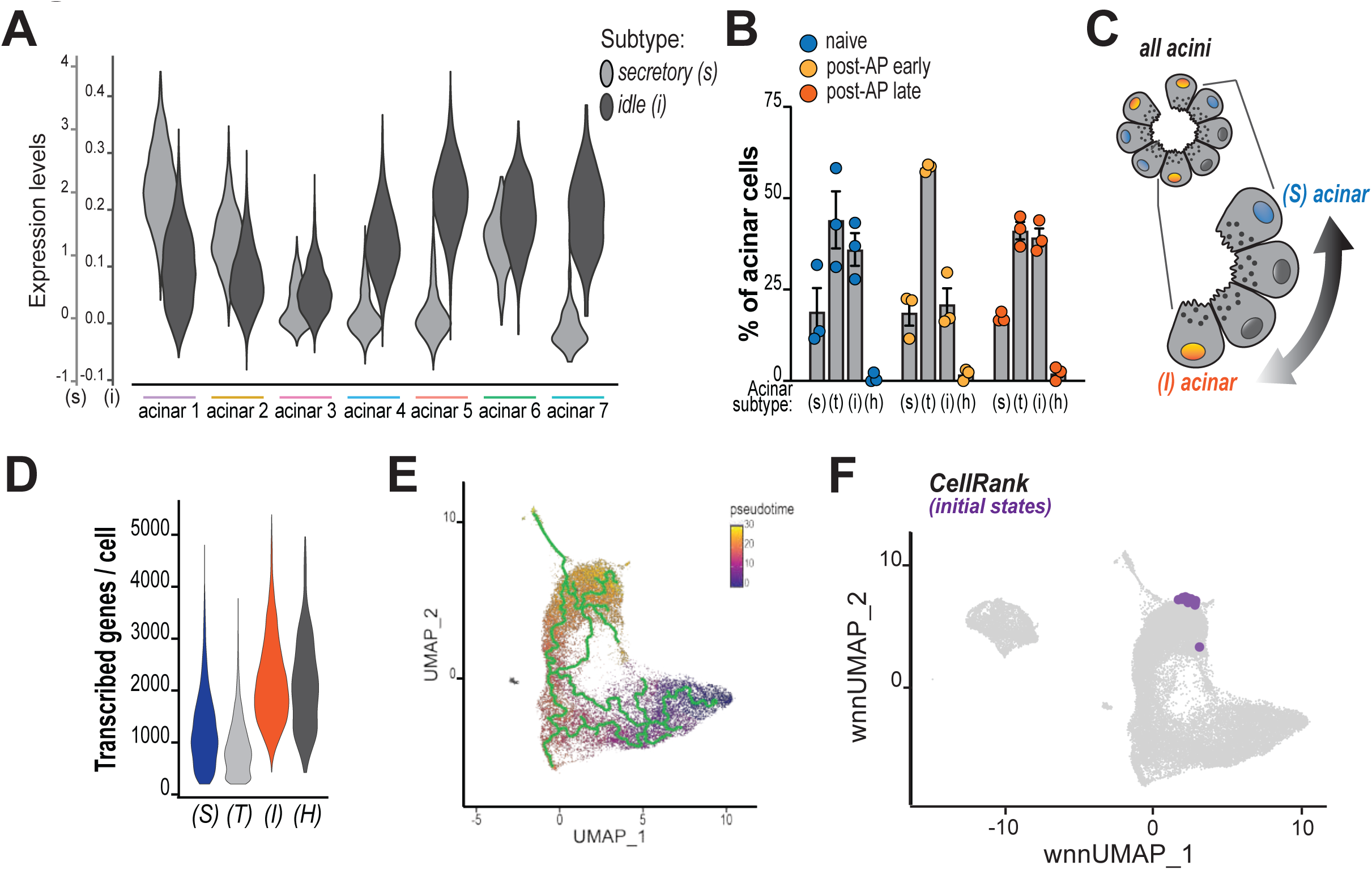
secretory and idling acini are phenotypically plastic. **A**, Violin plot showing the expression score of secretory and idling acinar signatures (light and dark gray, respectively) for all acinar subclusters. **B,** Relative abundance (expressed as percentage of all acinar nuclei) of secretory (S), transitional (T), idling (I) and hybrid (H) acinar metaclusters at all time points. Each dot denotes a mouse. **C,** schematic representation of the phenotypical spectrum of acinar specification. Gray nuclei indicate transitional acini. **D,** Violin plot showing the number of expressed genes per cell in each acinar metacluster. **E,** pseudotime trajectory designed by Monocle3 and highlighted by the green line. Starting point was assigned based on the cluster with the most cells in naïve mice. In **F,** wnnUMAP of acinar and ductal nuclei (light gray dots) as in Figure 2C superimposed with purple dots that denote CellRank2-predicted initial states (probability) along computed trajectory.

Our integrative analysis enabled the definition of cellular states that lie intermediate within a phenotypical spectrum that is defined by secretory and idling populations at the extremities (Figure 5), the former characterized by a more canonical exocrine signature and heightened expression of secretory genes. In contrast, idling cells are characterized by a more variegated transcriptome (Figure 5D), which includes proteins dedicated to signal transduction and overlaps. In addition, idling acini exhibited features observed in pancreatic epithelial cells during AP (Suppl. Figure 5C)^25,43^. Within this spectrum, we posited that transitional (*T*) acini might be incidental to phenotypic transition across plastic acinar cell states. Indeed, pseudotime reconstruction showed that clusters “1” and “5” are edge points at opposite ends of a transcriptional spectrum (Figure 5D), and were connected by a poorly defined trajectory that runs across clusters “2”, “3” and “4”, which corroborated their status of transitional cells. Of note, (*T*) acini were characterized by higher levels of the progenitor-associated marker *Cd44*^44^ (Suppl. Figure 5D).

A qualitative assessment of pseudotime trajectory suggested that (*S*) cells were at the dead end of phenotypic transition. In contrast, (*I*) acini denoted the ability to further evolve into more complex states, including proliferative acinar cells and the phenotypically hybrid (*H*) cluster “6” (Figure 5D). This further aligns with entropic analysis of chromatin/transcription connection (Figure 4D), which suggest that cluster 5 (which represents >95% of idling acini) is endowed with superior plasticity. To corroborate this hypothesis, we applied CellRank analysis^26^, which integrates multiple analytical kernels to identify APEX states in cell fate mapping. We found that idling acinar cells clearly constitute initial states (Figure 5F), which can be defined as apical (or loosely, more plastic) populations in cell fate trajectories. Initial states overlapped with areas of elevated AP-1 activity (Suppl Figure 5E), which has been linked to acinar lineage infidelity during AP-induced carcinogenesis^2,25,27^. Finally, this is consistent with pseudotime trajectory that highlights the idling metacluster as a focal point around which cells pivot toward different fates and point to this acinar subset as a regulatory node for acinar cell dynamics.

### Idling acinar cells exhibit plasticity during pancreatic regeneration

We next examined the behavior of acinar idling cells in the recovery from AP. In line with their superior sensitivity to environmental stimuli^13^, their abundance declines early after AP concurrent with increase in transitional, hybrid and proliferating acinar cells (Figure 6A). We speculated that all these states originate from idling acini that are positioned in a more apical transcriptional/epigenomic conformation (Figure 5F). To this goal, we asked whether idling-like features are required for pancreatic regeneration and expand in post-AP acini. We first interrogated chromatin elements that determine acinar cell heterogeneity and found 1723 regions that exhibited differential accessibility in (*I*) cells - compared to (*S*). Interestingly, loci associated with neoplastic progression^25^ were more accessible in idling cells (Suppl. Figure 6A) albeit overlap with profiles generated from FACS-sorted cells was minimal (Suppl. Figure 2C). We developed an “acinar (*I*) chromatin signature” and mapped the accessibility of those loci in naïve and post-AP pancreata to find that the makeup of idling cells propagates to (*T*), (*H*) cells early after pancreatitis and eventually contaminates also a fraction of (*S*) cells (Figure 6B).

**Figure 6.**
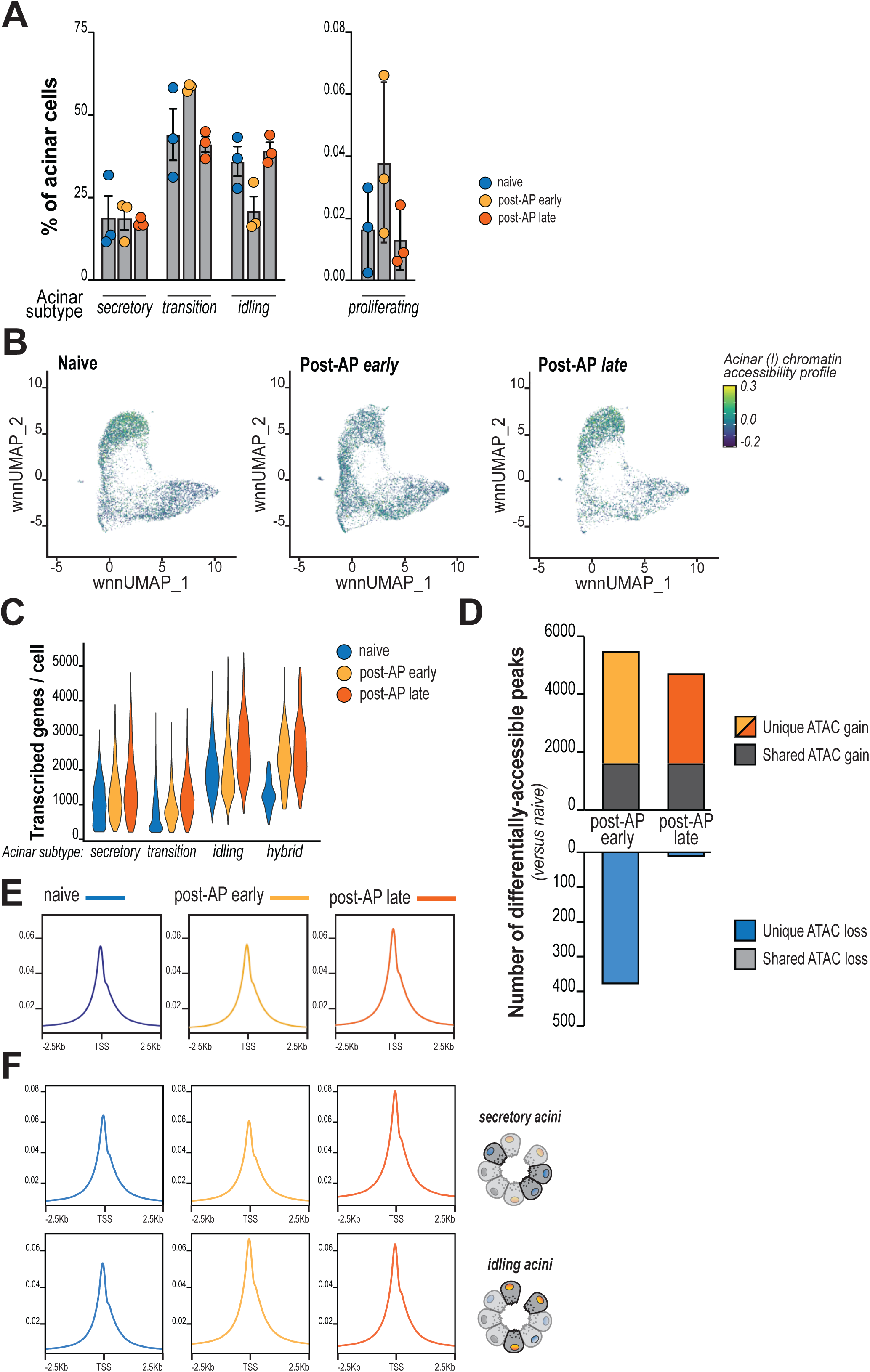
secretory-to-idling chromatin switch contributes to epigenetic reprogramming after AP. **A**, relative abundance of acinar metaclusters and proliferating cells across different timepoints pre- and after AP. **B,** Feature plot showing distribution and levels of idling cell chromatin signature across different timepoints pre- and after AP. **C,** Violin plot showing the number of expressed genes per cell in each acinar metacluster across different timepoints pre- and after AP. **D,** Number of ATAC peaks that significantly gained (top) or lost (bottom) accessibility in post-AP compared to naïve pancreas. Bar color indicates experimental condition and/or ATAC gain vs loss. Peaks that gained (or lost, =0) accessibility in both indicated conditions were highlighted in dark gray. Peaks were computed from *pseudobulked* acinar cells. **E-F,** Metagene representation of the mean ATAC-seq signal for regions around the transcription start site (TSS) of acinar cell-expressed genes. Conditions are indicated in the legend (top) and color-coded. **F** shows metagene coverage for secretory (top panels) and idling (bottom) acini, separately.

Incidentally, transcriptome complexity also augments in (*T*) and (*H*) cells at 7 days post-AP and is elevated in all acinar subtypes at 28 days (Figure 6C). Although evidence is correlative, these data suggest that acinar idling cells are most significantly perturbed by AP and react undergoing fission or a partial phenotypic skewing toward transitional or – to a lesser extend – hybrid states. Yet, their chromatin makeup is conserved across cell plasticity and determines increased complexity of the whole acinar transcriptome.

### Tissue regeneration induces permanent chromatin opening at acinar cells

To examine durable consequences of tissue homeostasis, we mapped chromatin accessibility in naïve and post-AP acinar cells using a *pseudobulking* approach. As expected, accessible peaks are invariably associated to actively transcribed genes (ca. 90% of detected ATAC peaks, irrespective of condition) and proportionally located within gene bodies, promoters and enhancers (Suppl. Figure 6B-C). We found 13010, 810 differentially-accessible peaks (DAPs) in post-AP *early* and *late* acini, respectively. For the most part, DAPs localize at enhancers and are clearly enriched in AP-1 binding motifs (Suppl. Figure 6D), but no significant enrichment for specific gene ontology domains could be found. DAPs were not linked to differential expression of proximal genes (Suppl. Figure 6E-F) that together indicates indirect impact on active transcriptional network.

Remarkably, nearly all DAPs in post-AP *late* samples were associated to gains in chromatin accessibility, the large majority occurring at idling and transitional acini (Figure 6D; Suppl. Figure 6G). In contrast, post-AP *early* acini exhibited more heterogeneous changes, with only a fraction of those ultimately conserved in post-AP *late* cells. These findings document widespread and permanent opening of acinar cell chromatin after pancreatitis, in line with previous studies^2,3^. They also indicate a progressive remodeling of the epigenome that evolves long after the resolution of local inflammation and possibly linked to phenotypic adaptations of histological identical acinar cells after recovery from pancreatitis. This was also evident when profiling chromatin accessibility around transcription start sites (TSS) that highlights increase in chromatin opening between 7 and 28 days (Figure 6E; also observed at enhancers: Suppl. Figure 6H).

We speculated that subpopulation dynamics could explain non-linear evolution of chromatin accessibility. To tackle this hypothesis, we characterized epigenetic differences of secretory and idling acinar cells. Time-point examination demonstrated early (7 days) and steady (conserved at 28 days) opening of TSS after AP for idling acinar cells, while surprisingly, secretory cells showed marked increase in accessibility only in post-AP *late* cells (Figure 6F). This aligns with the concept that idling acini are primarily sensitive to environmental cues; at the same time however, our data demonstrate that inflammation-induced chromatin opening eventually propagates to all acinar subtypes. The timing (early to late post-AP) could indicate that proliferation of idling acini does not contribute to pervasive epigenetic reprogramming. We speculated that AP induces a silent phenotypic transition, where secretory acini acquire an epigenetic makeup similar to idling cells. This is in part corroborated by our observations. First, pseudotime trajectory shows that *a)* post-AP acini evolve toward clusters “5”, “6” and “7”, *b)* phenotypic skewing can occur through at least two different routes (via “transition” or via “hybrid” acini) (Figure 5E). In addition, transitional and secretory acini showed enhanced chromatin accessibility at neoplasia-associated loci that typify idling acinar cells (Suppl. Figure 6I). Ultimately, post-AP *late* acini showed skewing toward an idling-like chromatin profile, with the majority of acinar cells acquiring a more diversified transcriptome (Suppl. Figure 6J).

### Population dynamics and chromatin opening is associated to permanent reprogramming of post-AP acinar cells

To assess consequences for cell functions, we examined RNA expression levels from all 23501 acinar cells to detect 10014 expressed genes and found extensive reprogramming post-AP; of note, “*early*” and “*late*” acinar cells upregulate distinct sets of genes (Figure 7A, shows top upregulated genes per condition). However, acinar cells exhibit marked transcriptional differences at 7 days, but their transcriptomic profile largely overlaps with naïve cells at 28 days post-AP (Suppl. Figure 7A). This again demonstrates that the exocrine pancreas progressively evolves after pancreatitis despite the termination of histological remodeling.

**Figure 7:**
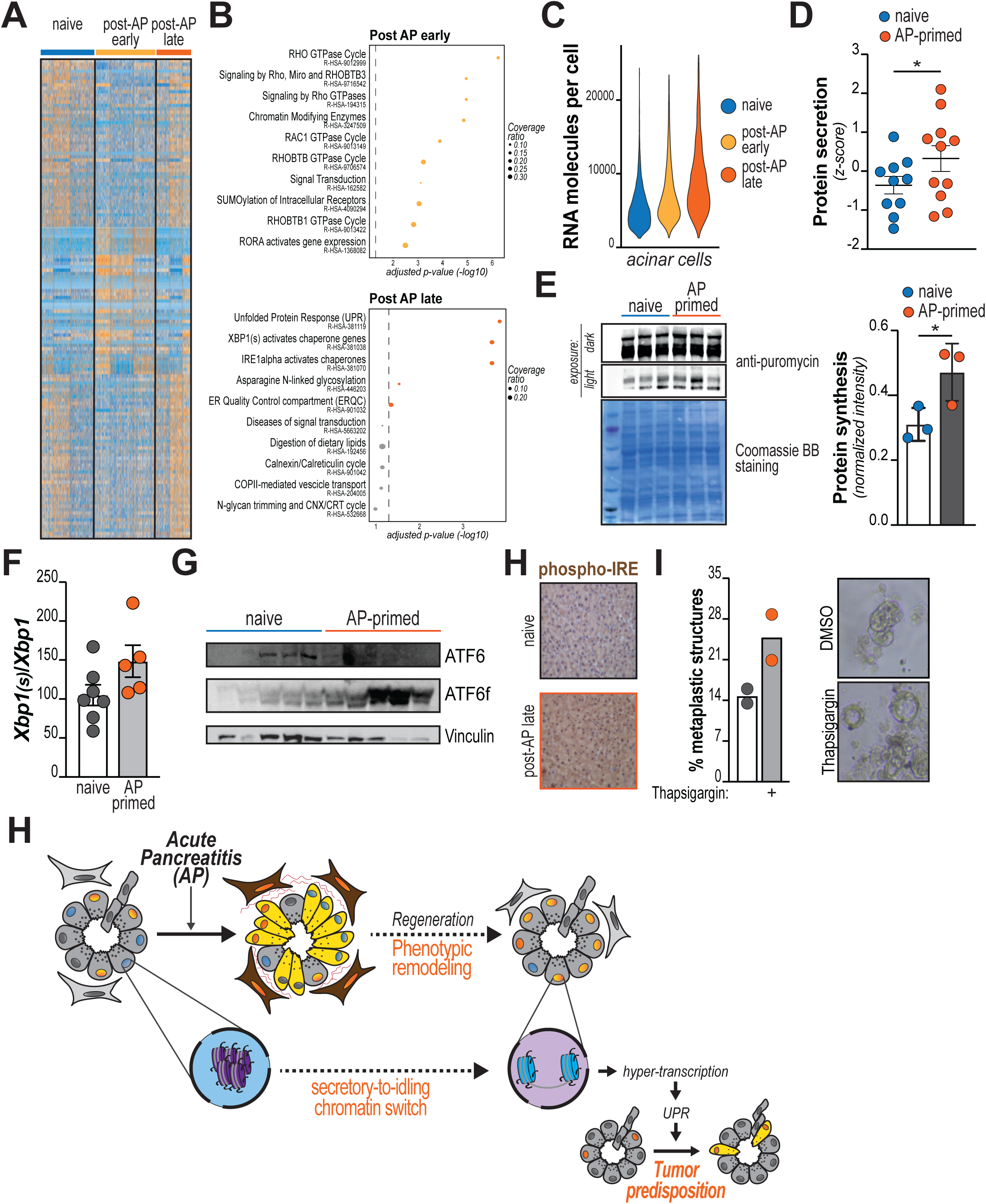
hypertranscription-associated UPR stress promotes metaplasia. **A**, Heatmap showing top 500 up-regulated genes for all indicated conditions in *pseudobulked* acinar cells. **B,** gene ontology analysis computed for all up-regulated genes at indicated conditions. Statistically significant associations were highlighted by colors that denote experimental conditions. Dashed lines mark *p*-value of 0.05. **C,** RNA molecules (UMIs) per nucleus in indicated samples. **D,** acinar explants were isolated from naïve (saline-injected) or AP-primed (cerulein-injected, 28 days recovery) pancreata and cultured in Waymouth’s medium. Both secreted (in the culture broth) and intracellular proteins were purified and quantified (*see Methods*). Each dot denotes a mouse. **E-H,** saline- or cerulein-injected mice were allowed to recover for 28 days (naïve and AP-primed, respectively; *n=3* mice/condition). In **E,** mice were administered puromycin via i.p. injection before sacrifice. Whole pancreas proteins were recovered and puromycin incorporation revealed by Western Blotting (left panel). On the right, quantification (ImageJ). In **F,** RNA was purified from whole pancreata and relative levels of *Xbp1* splicing quantified as ratio of spliced:unspliced transcripts, which were individually quantified for each sample. Bars show mean of biological replicates ± S.D. for indicated conditions. Each dot denotes a mouse. In G, whole pancreas proteins were recovered and levels of cleaved ATF6 highlighted. In **H,** pancreata were formalin-fixed and embedded in paraffin for immuno-histological analysis. Shown are representative images for immunohistochemistry (IHC) against phosphorylated IRE1. **I,** acinar explants were purified and embedded in Matrigel®. Upon solidification, spheroids were immediately treated with either Thapsigargin (50 ng/mL) or vehicle (DMSO) for two days. At end point, cytological images were acquired and examined for the presence of duct-like structures (*n=3* mice/group; *>30* images each prep). Shown are quantification of duct-like structures per optical field (values are percentage of all cell clusters examined; bars show mean of two biological replicates). **J,** schematic representation of working model. For all panels, columns show mean, ± SD (*, *P* < 0.05; **, *P* < 0.01; ***, *P* < 0.001)

Gene ontology analysis showed that top upregulated genes in acinar cells from “post-AP early” mice are highly enriched in members of the RHO GTPase family (Figure 7B). These proteins are essential for the remodeling of cytoskeleton and play an important role in acinar metaplasia^45^. RHO-dependent signaling is elevated in idling cells and enhanced activity might support regeneration (Suppl. Figure 7B-C).

Immunofluorescent staining confirmed the presence of actin stress fibers in pancreata of cerulein-treated mice but only at the earlier time point (Suppl. Figure 7D). These structures are elicited by RHO mechanosensing as demonstrated by increased phosphorylation of Myosin Light Chain-s (MLC2), a protein involved in the transduction of mechanical cues (Suppl. Figure 7E). This is in line with the idea that RHO^high^-idling acinar cells colonize most of the regenerating pancreas after the restoration of tissue architecture, likely exploiting phenotypical promiscuity of acinar cell states. This also leads to durable impairment of acinar cells’ mechanosensing. We asked whether downstream effectors might exacerbate acinar cells plasticity and predispose to tumor initiation. Indeed, we found the targeting mechano-transduction with fasudil (ROCKi) + MLCK inhibitor suppressed metaplasia of primary acinar explants *ex vivo* (Suppl. Figure 7F) although seemingly promoted cell proliferation. These findings demonstrates that AP induces- and acinar cells retain impairment in mechanosensing.

### Elevated chromatin accessibility promotes UPR to predispose for oncogenesis

We anticipated that the spreading of an idling-like chromatin profile leads to enhanced transcriptomic complexity in all pancreatic acinar cells. Consistent with a widespread and long-lasting increase of chromatin opening, RNA transcripts were markedly increased in post-AP *late* acini (Figure 7C), indicative of hyper-transcription. A more granular examination showed that the count of RNA molecules per cell increases significantly in hybrid and transitional cells (Suppl. Figure 8A), which more directly originate from idling cells and exhibit elevated transcription from baseline. Secretory cells more modestly increase their transcriptional outputs over time (Suppl. Figure 8A). In striking contrast, the number of UCI per ductal cell was not affected (Suppl. Figure 8B).

We sought to understand the mechanistic link between tumor susceptibility and heightened transcription and speculated this may elicit proteostatic and/or synthetic stress, which has been associated to tumor initiation^43^. In line, we found that AP-primed acinar explants showed elevated protein synthesis and secretory activity (Figure 7D-E), which are commonly associated to adaptive Unfolded Protein Response pathway (UPR). RNA expression analysis showed that UPR-associated genes are persistently upregulated in post-AP late acini (Figure 7B). In addition, we examined cerulein-injected mice and found consistent elevation of UPR markers in inflammation-primed pancreata, including increase of spliced *Xbp1* transcripts, IRE1 phosphorylation and ATF6 cleavege (Figure 7F-H). Finally, we observed that AP-sensitive idling acini were characterized by chronically-elevated UPR (Suppl. Figure 8C-D), consistent with the notion that idling-like features propagates to the entire exocrine compartment.

These findings suggest that sustained UPR signaling is a mark permanently imprinted on acinar cells by local inflammatory events and might mediate elevated oncogenic susceptibility. To tackle this point, we treated Matrigel-embedded primary acinar cells with sublethal doses of UPR inducer Thapsigargin and performed morphological examination after 2 days. We found that Thapsigargin induced a marked increase in the number of duct-like structures (Figure 7I), confirming that moderate ER stress facilitates acinar cell metaplasia (also seen in REF:^43^).

Interestingly, induction of UPR stress by Thapsigargin did not further enhance metaplastic potential of AP-primed acinar cells (Suppl. Figure 8E) to indicate that pancreatitis does not sensitize acinar cells to exogenous stimuli but rather sustains a long-lasting cell-intrinsic response.

## Discussion

We applied single nucleus technology for the paired mapping of cellular transcriptome and chromatin accessibility of cells in the pancreatic tissue. Those were documented in cohort of mice with no histological, structural or functional differences, but different history of local inflammation. Together our data indicate that a single episode of pancreatitis induces a complex homeostatic response, which includes phenotypical remodeling of exocrine cells and pivots around a subpopulation of “idling” acinar cells. The process leads to progressive and indiscriminate opening of acinar cells’ epigenome, augmented synthetic activity and chronic proteostatic stress, which in turn facilitates acinar-to-ductal metaplasia (Figure 7H). These results mechanistically tie acute pancreatitis, tissue dyshomeostasis, epigenetic reprogramming and tumor predisposition.

Memory of inflammation is well characterized in innate immune cells, where is referred to “immunological training”^46^. Skin and intestine stem cells also develop a similar form of memory in response to proximal or distant inflammation^47,48^. The pancreas lacks classical stem cells^7^ yet epithelial cells are permanently primed by experimental pancreatitis^2,3^. Because acinar cells exhibit actionable plasticity, it was plausible that cell de-differentiation enables the establishment of epigenetic memory at specific – *i.e.*, amenable – cell states. Indeed, co-option of transient cellular exocrine states has already been described^27^, while inflammation clearly impact the epigenome^21,30,49^. In this respect, our study is the first comprehensive description of chromatin reprogramming across phenotypic states.

The epigenetic nature of inflammation memory has been previously described by others using bulk cell approaches from lineage-traced, FACS-sorted exocrine cells^2^ or *in vitro* ductal organoids^3^. While these studies are keystones in our understanding of epigenetic memory triggered by acute pancreatitis, our work is able to integrate population dynamics and analyze the epigenetic reprogramming of acinar cells in the context of their phenotypic plasticity. Indeed, we found that inflammation alters the epigenome of “idling” acinar cells, a subset characterized by elevated reactivity to external stimuli^13^. In line with prior work^2^, we found that the epigenetic memory of pancreatitis progressively evolves for several weeks after AP induction/resolution. We were able to link this evolution to a spreading “secretory-to-idling” epigenetic switch, where more canonical “secretory” acini acquire a chromatin configuration similar to “idling” cells.

Also similar to prior work^2,3^, we found durable increase of chromatin accessibility after pancreatitis, with open loci associated to AP-1-binding sites. However, our analyses indicate that AP-1 is most frequently recruited at either enhancers or non gene-associated sites (*data not shown*). This aligns with the notion that AP-1 family members could dictate enhancer selection and possibly function as pioneer factors^50–52^, rather than only mediate acute stress responses.

Our data suggest that acinar cells are exposed to chronic UPR stress after memory is established. This facilitates ADM (Figure 7I) and carcinogenesis upon oncogene expression^43^. We moved to identify the source of UPR stress, as AP-primed acini exhibit elevated transcriptional and synthetic capabilities, with no obvious selectivity. This might be linked to undiscriminated chromatin opening observed after AP and aligns to the notion of hypertranscription^53^. In fact, augmented transcription is generally pro-oncogenic, as exemplified by widespread activity of c-MYC and by the anti-cancer properties of bromodomain inhibitors^54^. Chromatin remodeling-induced hypertranscription supports pancreatic cancer progression in mice^54^, but how generalized expression initiates carcinogenesis was still an open question. Acini are synthetic cells, specialized in the production and secretion of a limited number of proteins. A more variegated proteome – like the one of “idling” acini – correlates with the establishment of a mild proteostatic stress. Our experimentation links phenotypical switch of acinar cells, to adaptive UPR stress and tumor initiation. It will be interesting to assess whether UPR-alleviating interventions can reduce cancer susceptibility in AP-primed mice.

Besides biological findings, our work incorporates technological and computational advances. First, we profiled paired transcriptomic and epigenomic profiles from individual nuclei. This refines the analysis of hard-to-dissociate tissues – like the pancreas. Because we harvested cells from label-free, frozen pancreata, the methodology we employed avoids the introduction of mechanical and biological biases (*i.e.*: expression of exogenous genes, cell selection, protease digestion, cell dissociation, FACS sorting) that can significantly impact the read-out. Next, we could identify and analyze all expected cell populations in the pancreas, at logical abundance.

While this had always been challenging for pancreas studies, it allowed us to examine effects induced by AP in a holistic manner. We investigated alterations in near all cell types identified (Figure 3 and *data not shown*) and only focused on acinar cells after applying Bayesian statistics across all cell populations.

Finally, we built upon the concept that changes in chromatin accessibility might poise for transcriptional activation and prime pancreatic epithelial cells for plasticity^26^. The entropic correlation between unpaired transcriptomic and epigenomic profiles was previously calculated through the generation of metacells^26^, a compelling but reductionist approach. We were able – for the first time – to calculate entropy in the transcription/epigenome relationship at single nucleus resolution, in a fully conservative approach that corroborates the plastic identity of pancreatic acinar cells and unveiled the propensity of “idling” and “hybrid” acini to transdifferentiate to ductal-like cells, enticingly pointing at specific acinar phenotypical subsets as tumor-predisposing states.

To conclude, we characterized how inflammation memory is established in the pancreatic parenchyma. We also found that AP induces an hypertranscription phenotype in acinar cells that activates an adaptive UPR response. Our data identify protein dyshomeostasis as a targetable determinant of pancreatic cancer predisposition.

## Supporting information

Suppl. Figure altogether

## Acknowledgments

This project has received funding from the European Union’s *Horizon 2020 research and innovation programme* under grant agreement No. 824110. A.C. thanks Steven Zhao for discussing the project’s idea at its inception and convincing on the power of single cell technology for the project. A.C. and M.F. also thank Giuseppe Diaferia and Pierluigi Di Chiaro for helpful discussions and manuscript editing. A.C. acknowledges funding from the Italian Association for Cancer Research (AIRC; ref: MFAG-#23029). M.F. thanks IBSA Foundation for salary support.

## Author contributions

M.F., G.F. performed experiments. M.F. coordinated bioinformatic work and generated all pipelines. M.F, J.L., S.L., V.I. all performed bioinformatic analyses. J.L. prepared sequencing libraries and performed NovaSeq runs. A.C. and M.F. designed all figures. C.Ca. provided histological samples. A.C. and C.Co. provided funding support, conceptualized and supervised the work, wrote the manuscript.

## Materials and Methods

### Transmission electron microscopy

Tissue samples were harvested and fixed for one hour at 4 °C in 2.5% glutaraldehyde (Sigma-Aldrich – G5882) in sodium cacodylate buffer 0.1M pH 7.4 (Sigma-Aldrich - C0250) and washed using sodium cacodylate 0.1M pH 7.4. Cells were incubated with osmium tetroxide 1% (Sigma-Aldrich – 201030) and potassium ferrocyanide 1% (Sigma-Aldrich – 455989) in sodium cacodylate buffer 0.1M pH 7.4 for 1h at 4° C, embedded in epoxy resin and sectioned. Ultrafine (60-80 nm) dissections were performed with Ultratome V (Leica) ultramicrotome and contrasted with 1% of uranyl acetate (Electron Microscopy Sciences – 22400) and 1% of lead citrate (Electron Microscopy Sciences – 17800). Thin sections were imaged on a transmission electron microscope FEI Tecnai G2 with OSIS Veleta cameras and visualized at a magnification of 37000 X.

### Acinar cell isolation and metaplasia

Acinar cells were isolated as previously described. Briefly, pancreata from mice were collected upon sacrifice, washed twice in cold Hank’s Balanced Salt Solution (HBSS, Biowest - L0611) and subsequently minced with a sterile razor blade. Tissue was then digested with 1 mg of collagenase P (Roche - 11213865001) in 5 mL HBSS at 37°C for 21 minutes in agitation and interrupting the digestion every 7 minutes to mechanically disrupt the clogs by pipetting with progressively smaller pipette tubes. The reaction was stopped and tissue homogenate was washed twice with HBSS containing 5% calf serum (CS, Biowest - S0400) and then filtered through a 500 μm steel mesh and a 100 μm cell strainer. The flowthrough was carefully laid onto an HBSS +30% CS solution and centrifuged at 200g for 5 minutes in order to pellet only multicellular acinar structures.

To evaluate the duct-forming capabilities of acinar cells as a proxy of metaplastic capabilities, isolated acinar clusters were resuspended in a 1:3 mix of Matrigel-GF reduced (Corning – 356231) and Waymouth’s medium (Sigma-Aldrich – W1625), containing 10% CS, 0.1 mg/mL trypsin inhibitor (Sigma-Aldrich – T2011) and 1% penicillin/streptomycin (Biowest - L0022).

Suspension (150 μL) was seeded onto a 48-well plate and incubated at 37°C for solidification for at least 1 hour. Upon Matrigel solidification, 300 μL of Waymouth’s medium, supplemented as previously described, was added to each well. Cells were monitored daily, and images were acquired at day 2 using a Nexcope NIB620 inverted optic microscope. At least 20 HPF were quantified in single blind conditions by an independent operator for each experimental time point.

For differential stiffness plating, isolated acinar clusters were resuspended in a stiff matrix composed by 1:1:1 mix of Matrigel-GF reduced : rat collagen I neutralized solution 4.43 ng/mL (R&D Systems – 3440-100-01) : cell suspension in complete media or in a matching soft matrix in which collagen part was substituted with saline solution. Then, the rest of the experiment was performed as aforementioned.

In case of treatment supplementation, drugs have been added both in the 3D matrix and in the above supernatant at the indicated final concentration. Treatment employed were thapsigargin (Sigma-Aldrich - T7458), fasudil (Sigma-Aldrich - CDS021620).

### Fecal protein content quantification

To assess total fecal protein content, feces were collected, weighted and resuspended in 1 mL of fecal lysis buffer (2% SDS, 150 mM NaCl, 0.5 mM EDTA), sonicated in an ultrasonic bath at full power for 30’, resuspended with a plastic Pasteur pipette and centrifuged 5’ at 13000g to remove the particulated suspension. Protein concentration in the remaining supernatant was assessed using a Bradford reagent assay according to manufacturer’s protocol (Sigma-Aldrich - B6916), and normalized on collected fecal weight and on animal weight.

### Blood glucose measurement

Blood glucose levels were measured from tail of animals prior to sacrifice using a portable glucometer (Sinocare Safe AQ Smart).

### Histological analysis of murine pancreatic tissue

For histologic evaluation, tissue samples were harvested as described^30^. After pancreas excision, the organ was laid on a planar surface and fixed with formalin overnight. Sectioning of paraffin-embedded tissues exposed the transverse axis and revealed whole organ morphology (4 μm sections).

For hematoxylin and eosin staining, tissue sections were dewaxed and rehydrated. Hematoxylin staining was applied for 3 minutes to color cell nuclei, while eosin cytoplasmic counterstain was subsequently employed for 1 minute. After dehydration, sections were mounted with a xylene-based mounting medium and imaged using a Nikon Eclipse Ci-S optical microscope.

For immunohistological analysis, tissue sections were dewaxed and rehydrated. Antigen retrieval was performed by boiling samples in citrate buffer for 20 minutes and endogenous peroxidase was blunted by incubating samples with 3% H2O2 for 10 minutes. After blocking with a 2% bovine serum albumin solution for 30 minutes, primary antibody was incubated overnight at 4°C. Immunohistochemical coloration was performed using a commercial kit (Histoline – MTM001) according to manufacturer instructions, and nuclear counterstain was obtained via incubation for 15 minutes with hematoxylin. After dehydration, sections were mounted with a xylene-based mounting medium and imaged using a Nikon Eclipse Ci-S optical microscope. For each animal, at least 20 HPF were imaged and quantified; number of animals employed is stated in each figure legend. Representative images are shown in figures.

For PicroSirius Red staining, tissue sections were dewaxed and rehydrated. After these steps, tissue section was covered with Sirius Red F3B 0.1% (Sigma-Aldrich – 365548) in a saturated picric acid solution (Sigma-Aldrich – 197378) for 1 hour. Following incubation, coloration was differentiated by dipping 10 times in two different baths of acetic acid 0.5% (Sigma-Aldrich – 33209). Imaging was performed using a Nikon Eclipse Ci-S optical microscope.

### Immunofluorescence staining of murine pancreatic tissue

Tissue samples were harvested as described^30^. After pancreas excision, the organ was laid on a planar surface and fixed with formalin overnight. Sectioning of paraffin-embedded tissues exposed the transverse axis and revealed whole organ morphology (4 μm sections). Tissue sections were then dewaxed and rehydrated. Antigen retrieval was performed by boiling samples in citrate buffer for 20 minutes. After blocking with a 3% goat serum (Sigma-Aldrich – S26) solution in PBS for 1 hour, primary antibody was incubated overnight at 4°C. Following, sections were incubated with matching host secondary antibody conjugated with the indicated fluorochrome for 1h at room temperature. Nuclei were counterstained with 0.01% Hoechst 33258 (Life Technologies – H3569) for 5 minutes, and sections were mounted using ProLong Glass mounting medium (Thermo Fisher Scientific – P36982).

Imaging was performed using a Zeiss LSM900 inverted confocal microscope equipped with a Airyscan2 detector; at least 20 images for each experimental point were acquired.

### Nuclei isolation and library generation for single-nuclei multiomics

Pancreata from mice were collected upon sacrifice; after collection, first, a small piece of each clamp frozen pancreas piece was pulverized by using a liquid nitrogen cooled Cellcrusher.

Nuclei were isolated from tissue powder employing a protocol for isolating nuclei from RNAse-rich tissue that uses citric acid, previously described^13^.

In short, one small spoon of tissue powder (5-10 mg approximately) was suspended into 1 mL of pre-cooled S25 buffer, composed by 1.5M sucrose (Sigma-Aldrich – S7903), 250mM citric acid (Sigma-Aldrich – C0759), 0.1% Hoechst 33258 (Life Technologies – H3569), inside a glass douncer tissue grinder (DWK Life Sciences - 357358). With 5 strokes of a tight pestle every 3 minutes for 10 minutes, nuclei were released into the buffer. Subsequently, the lysate was transferred through a 35 µm strainer into a 1.5 mL Eppendorf tube and centrifuged at 4°C for 5 min at 500 g. Afterwards, the supernatant was removed and the pellet was resuspended in 500 µL S25 buffer. To remove clearly visible cell debris, a density centrifugation was performed, by carefully adding 500 µL S88 buffer, composed by 1.5M sucrose and 250mM citric acid, to the bottom of the Eppendorf tube followed by centrifugation at 4°C for 10 min at 1000 g. After removal of the supernatant, the nuclei were resuspended in 250 µL S25 buffer, their integrity was checked and they were counted with a CountessTM II FL Automated Cell Counter (Thermo Fisher Scientific). The samples were spun down again at 4°C for 5 min at 500 g, filtered through a 10 µm strainer and resuspended in 100µL resuspension buffer.

The nuclei suspension was diluted to load 10000 nuclei per sample onto the 10x Genomics Chromium controller. Libraries were generated with the Multiome Kit (10x Genomics - PN-1000283) following the 10x Genomics Chromium Next GEM Single Cell Multiome ATAC + Gene Expression user guide and the finished libraries were sequenced on an Illumina NovaSeq6000 (S4, paired-end). The sequencing data is deposited on the European Nucleotide Archive under the accession number PRJEB57065.

### Pre-processing and quality control of sequencing reads

The sequencing data was aligned to the mouse genome (mm10/GRCm38) using cellranger-arc (10x Genomic - v2.0.0) and afterwards aggregated. To reduce the noise caused by ambient RNA, the R package SoupX2 v1.5.2^55^ was used with the manually set gene sets described in the table below.

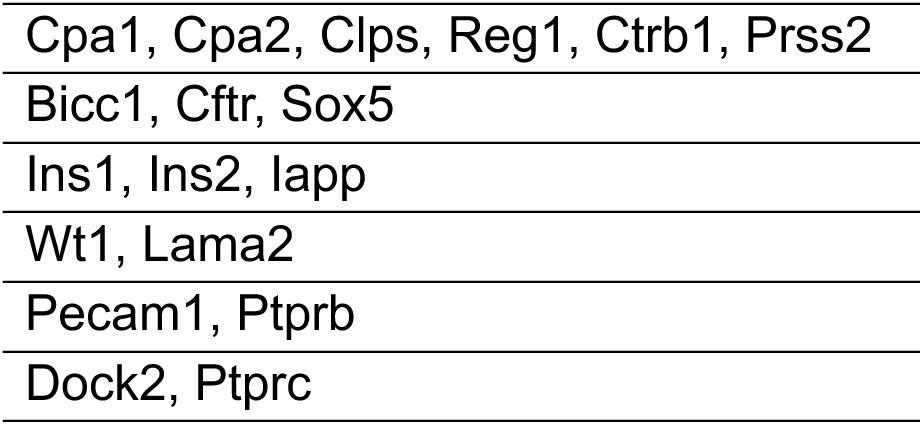

The data was filtered for quality control, subjected to unsupervised clustering, integrated and analyzed using the R packages Seurat v4.1.0^31^ and Signac v1.6.0^56^. During quality control all nuclei with less than 100 genes and 1000 molecules and more than 30,000 molecules in the RNA assay, 160000 molecules in the ATAC assay and more than 5 % mitochondrial genes were removed. The dimensional reduction, clustering and integration was done on both assays independently. Afterwards, a weighted nearest neighbors (WNN) graph was calculated combining the RNA and ATAC modalities.

### UMAP reduction and nuclei clustering

Upon WNN integration, Principal Component analysis was run on the integrated dataset employing the RunPCA command of the Seurat pipeline, and dimensions up to PC30 were selected using the ElbowPlot method to identify the top 3000 highly variable features using the FindVariableFeatures command.

Upon selection of variable features, neighbor identification and cluster selection was performed using respectively the FindNeighbors and FindClusters commands on the integrated WNN graph. Top 500 variable features for the RNA assay were employed to obtain an heatmap of expressed genes stratified for experimental timepoints using the heatmap base R function.

Cluster names were then assigned using a panel of well-described markers, listed in the table below:

| Cell type | Marker gene | Reference |
| --- | --- | --- |
| Acinar cells | Amy2a5 | Franzén, Gan, Björkegren, Database, 2019 |
|  | Ctrc |  |
|  | Cpa1 |  |
|  | Prss2 |  |
|  | Rbpjl |  |
|  | Try10 | Bult, Sternberg, Mamm Genome, 2023 |
| Proliferating acinar cells | Acinar markers | see above description of acinar cell markers |
|  | Mki67 | Bullwinkel et al., Journal of Cellular Physiology, 2005 |
| Ductal cells | Cftr | Marino et al., J. Clin. Invest., 1991 |
|  | Hnf1b | Reichert and Rustgi, J. Clin. Invest., 2011 |
|  | Onecut1 | Kropp et al., Cell. Mol. Gastroenterol. Hepatol. |
|  | Sox9 | Qadir et al., PNAS, 2020 |
|  | Krt19 |  |
| Intrapancreatic bile ductal cells | Positive for all ductal markers excluding Cftr | Hendley et al, eLife, 2021 |
| $\alpha$ cells | Gcg | DiGruccio et al, Mol Metab, 2016 |
|  | Ttr |  |
| $\beta$ cells | Ins1 | DiGruccio et al, Mol Metab, 2016 |
|  | Ins2 |  |
| $\gamma/\delta$ cells | Sst | DiGruccio et al, Mol Metab, 2016 |
|  | Ppy | Perez-Frances et al.,<br>Nature Communications,<br>2021 |
| Endothelial cells | Esam | Yokota et al., Blood, 2009 |
|  | Pecam1 | Müller et al., Experimental<br>and Molecular Pathology,<br>2002 |
|  | Vwf |  |
| Lymphatic endothelial cells | Flt4 | Kaipainen et al., PNAS,<br>1995 |
|  | Lyve1 | Jackson, APMIS, 2004 |
| Mesenchymal cells | Col1a1 | Hutton et al., Cancer Cell,<br>2021 |
|  | Msln | Khatri et al., Stem Cell<br>Research & Therapy, 2021 |
|  | Pdpn | Sunami et al., Cancers,<br>2020 |
|  | Vim |  |
| Activated fibroblasts | Fap | Elyada et al., Cancer<br>Discov., 2019 |
|  | Fn1 |  |
|  | Lama2 |  |
|  | Pdgfra |  |
| Pericytes | Cspg4 | Splunder et al., Trends in<br>Cell Biology, 2024 |
|  | Rgs5 |  |
| Adipocytes | Adipoq | Fitzgerald et al.,<br>Communications Biology,<br>2023 |
|  | Plin1 |  |
|  | Pparg |  |
| Glial cells | Mpz | Ghanavatinejad et al., J<br>Otol., 2019 |
|  | Ncam1 |  |
| Non-lymphoid immune cells | Cd68 | Holness and Simmons,<br>Blood, 1993 |
|  | Cfp | Schwaeble et al., Eur. J.<br>Biochem., 1994 |
|  | Csf3r | Touw and van de Geijn,<br>Frontiers in Bioscience,<br>2007 |
|  | Mrc1 | He et al., Cell Reports,<br>2021 |
| Lymphocytes | Ighd | Shi et al., Cell Discovery,<br>2019 |
|  | Ighm |  |
|  | Ltb | Stewart et al., Front.<br>Immunol., 2021 |
|  | Ptprc | de Koning et al., Front.<br>Immunol., 2021 |
|  | Trbc2 |  |

### Gene signature expression scoring

Gene signature expression was assigned to single nuclei in the analyzed dataset using the AddModuleScore function of the Seurat package.

Gene signatures used for the analysis are reported in Table S1.

### Quantification of cell population composition

To quantify differences in cell population composition in the multiomic dataset, cell subcluster amounts were embedded in a Python pandas dataframe and hierarchical information was superimposed during the preprocessing stage of the tascCODA pipeline v0.1.3^41^. tascCODA model was built with default settings, and iteratively performed using each of the cell subclusters as reference cell population. Hierarchical tree view was generated according to the pipeline, and only effect values classified as credible by tascCODA iterative procedure were plotted on the tree graph.

### Gene ontology enrichment analysis

Gene ontology enrichment was performed using the Reactome Pathway v2022 database^57^ through the Enrichr web portal^58^.

### RNA extraction and quantitative real-time PCR

Pancreata from mice were collected upon sacrifice; collected tissue were harvested in Qiazol lysis reagent (Qiagen - 79306) for total RNA extraction using phenol/chloroform organic extraction. Quantitative real-time PCR (qRT-PCR) analysis were carried out on cDNAs retrotranscribed with High Capacity cDNA RT kit (Qiagen - 4368814), and analyzed genes were amplified using 4x Capital qPCR Green Master mix (BiotechRabbit - BR0501702) on a QuantStudio 5 Real-Time PCR System (Thermo Fisher Scientific). For Xbp1 spliced and total quantification were performed in six different biological replicates, and each sample is the average of a technical duplicate. The quantification is based on the 2−ΔΔCt method on the average of the naïve samples.

### Pseudo-temporal ordering of single-nuclei

Single-nuclei RNA sequencing data were converted using the SeuratWrappers R package to a celldataset object suitable for further analysis. Preprocessing of exocrine cells data was conducted as described in the monocle3 pipeline^59^. Principal components, partitions and cluster assignments were manually transferred from the previously calculated based on the WNN graph obtained after integration. Then, pseudotime ordering and trajectory calculation was obtained using default settings with monocle3, and the starting root node of the resulting trajectory was chosen using the unbiased helper function described in the monocle3 vignette, choosing the node closer to the highest number of naïve acinar cells.

### Prediction of apex states with CellRank

Unbiased analysis of pseudo-temporal ordering of sequenced nuclei reproduces the putative chronology from naïve to early and late post-AP primed exocrine cells. However, substantial heterogeneity within each condition, particularly explained by the high degree of morphological and functional similarity of exocrine cells which has been reported by *in vivo* and *in vitro* experiments suggests that cells might be subjected by state transitions that cannot be defined by time series information alone. While this approach allows to describe transitions from a known origin^60^, complex alterations and fluxes between very similar populations often fail to be correctly described by a linear ordering along pseudo-temporal trajectories.

To avoid this issue, we resorted to use CellRank 2^61^ to be combine gene expression similarity with pseudo-temporal trajectories to robustly model the cell-state transitions as a Markov chain, successfully identifying initial apex states, inferred to be the sources of the cell-states observed in the data, describing putative hyperplastic cell populations.

To perform this analysis, we employed the Palantir package^60^ to compute pseudo-temporal ordering of exocrine cells starting from the initial node identified by aforementioned monocle3 analysis, and integrated it in a pseudotime kernel for CellRank2 analysis, as described in the package vignette. Moreover, following the default conditions described, we computed a pseudotemporal estimator based on these data, and applied the GPCCA algorithm to infer initial apex states.

### Correlation of exocrine subclusters features

To compute correlation between cell subclusters based on transcriptomic or epigenomic features, average expression of each feature was computed in the selected subgroups using the AverageExpression Seurat function, and then correlation maps were obtained by combining the gather function from the tidyr v.1.3.1 package with ggplot2 graphical geom_tile plotting^62^.

## Notes

### Competing Interest Statement

The authors have declared no competing interest.

