## Supplementary material for "Population dynamics after pancreatitis dictates long-lasting epigenetic reprogramming and mediates tumor predisposition": Suppl. Figure altogether

#### Supplementary Figures legends

**Suppl. Figure 1** (*related to Figure 1*). **A,** mice were in treated as described in main Figure. Before sacrifice, stool and blood drops were harvested. Proteins were extracted from stool samples and quantified (left panel), while blood glucose levels were measured with a standard glucometer (right panel). **B,** immunohistochemistry against Ki67 on pancreatic tissue sections. Three mice per group were analyzed and percentage of Ki67+ nuclei assessed for each optical field acquired. **C,** representative hematoxylyn and eosin staining of pancreatic tissue sections from saline- and cerulein-injected mice, sacrificed at indicated time points. **D,** representative immunohistochemistry (IHC) staining of pancreatic tissue sections against indicated leukocytic and endothelial markers from saline- and cerulein-injected mice, sacrificed at indicated time points. Quantification below.

**Suppl. Figure 2** (*related to Figure 2*). **A,** wnnUMAP plots of snRNA- and snATAC-sequencing (left and right panels, respectively. On the right, color coding of lineage-associated clusters. **B,** Relative abundance (expressed as % of total cells/nuclei analyzed) of cell populations obtained in our dataset, in Tosti's dataset and in a publicly-available single cell repository (TabulaMuris). **C,** RNA and ATAC peaks were overlapped with previously-published sequencing datasets obtained from FACS-sorted pancreatic epithelial cells (not perturbed, WT mice). **D,** ATAC-seq coverage tracks (color code as in C) for four representative cell populations (indicated) at loci associated to cell-identity genes. **E,** Feature plot showing the distribution of progenitor-like cells (indicated, in orange) within the acinar cell cluster. **F,** Feature plot showing the abundance of proliferating acinar cells in naïve (blue dots), post-AP early (yellow dots) and post-AP late (orange dots) pancreata. **G,** immunohistochemistry against Ki67 on pancreatic tissue sections from indicated mouse samples (n=3/group). Quantification on the right. **H,** dot plot showing the expression of markers of progenitor and metaplastic cells in proliferating and resting acinar cells. **I,** violin plots showing the expression score relative to gene signatures associated with pancreatic regeneration (Alonso-Curbelo et al, Nature, 2021) in proliferating and resting acinar cells.

**Suppl. Figure 3** (*related to Figure 3*). **A,** immunofluorescence staining of naïve mouse pancreata against Fibroblast Activation Protein (FAP, green) and alpha-Smooth Muscle Actin (aSMA, red). Nuclei were counterstained with DAPI (blue).

**Suppl. Figure 4** (*related to Figure 4*). **A,** relative abundance (expressed as percentage of all acinar cells analyzed) of cells assigned to indicated acinar clusters in in naïve (blue dots), post-AP early (yellow dots) and post-AP late (orange dots) pancreata. Asterisks denote statistically-significant changes.
B, correlation matrices showing vicinity of RNA (left) and ATAC (right) profiles among acinar cell subclusters. In the RNA matrix, orange lines define acinar metaclusters (secretory, transitional, idling and hybrid).

**Suppl. Figure 5** (*related to Figure 5*). **A,** violin plots showing the expression of secretory and idling acinar genes (left and right panels, respectively) in indicated metaclusters. **B,** wnnUMAP feature plots (restricted to the acinar cluster) showing imputed expression of secretory and idling acinar genes (left and right panels, respectively). **C,** Violin plots showing the expression of genes that characterize unperturbed murine acinar cells (N1, light gray) or cerulein-treated cells (N2, dark gray) in indicated metaclusters. **D,** wnnUMAP feature plots (restricted to the acinar cluster) showing imputed expression of *Cd44*.

**Suppl. Figure 6** (*related to Figure 6*). **A,** dot plot showing the overlap of sn-ATAC peaks for indicated metaclusters with ATAC peaks associated to pancreatic acinar cells (A1 and A2) or neoplastic epithelial cells (N1 and N2). **B,** bar graphs showing number of sn-ATAC peaks in acinar cells for condition and proportion of peaks associated to genes detected in sn-RNA-seq (i.e.: expressed in acini). **C,** bar graph and split pie chart showing the distribution of sn-ATAC peaks (acinar cells only). **D,** MEME analysis was performed on peaks that show different accessibility 28 days post AP. In orange, scores of heterodimeric members of the AP1 family. **E,** bar graphs showing number of differentially-accessible ATAC peaks (DAPs) in acinar cells and proportion of peaks associated to genes detected in sn-RNA-seq (i.e.: expressed in acini). **F,** bar graph showing the association of ATAC peaks or DAPs with expressed (gray boxes) or not-expressed (colored boxes) genes in acinar cells. **G,** proportion of DAPs in indicated acinar metaclusters (post-AP *late* vs naïve). **H,** waterfall plot showing chromatin opening at indicated conditions. **I,** dot plot showing the overlap of sn-ATAC peaks for indicated metaclusters with ATAC peaks associated to pancreatic acinar cells (A1 and A2) or neoplastic epithelial cells (N1 and N2).

**Suppl. Figure 7** (*related to Figure 7*). **A,** correlation matrix for acinar transcriptomic profiles at indicated conditions. **B,** Violin plots showing the expression of genes involved in RHO signaling in indicated metaclusters. **C,** Violin plots showing the expression of genes that characterize regeneration of thepancreatic epithelium in indicated metaclusters. **D,** representative images of immunofluorescence against ACTIN in pancreatic tissue sections. **E,** representative images of immunofluorescence against ACTIN and phospho-MLC2 (green and orange, respectively) in pancreatic tissue sections. Nuclei were counterstained with DAPI (blue). **F,** quantification of metaplastic structures (duct morphology) obtained from acinar explants treated with either a combination of ROCK and MLCK inhibitors (orange) or vehicle (gray). Each dot denotes an optical field.

**Suppl. Figure 8** (*related to Figure 7*). **A-B,** violin plots showing the number of RNA objects count per single nucleus in indicated cluster/metaclusters and conditions. **C,** wnnUMAP feature plots (restricted to the acinar cluster) showing imputed expression of genes involved in UPR or XBP1 signaling. **D,** Violin plots showing the expression of Xbp1 target genes in indicated acinar metacluster. **E,** quantification of metaplastic structures (duct morphology) obtained from acinar explants treated with Thapsigargin (50 $\mu$M). Acinar cells were purified from either saline-injected (gray) or AP-primed (orange) mice. Each dot denotes an optical field.

<u>For all figures</u>: genes signatures adopted in the study are listed in Suppl. Table 1

### Supplementary Figure 1

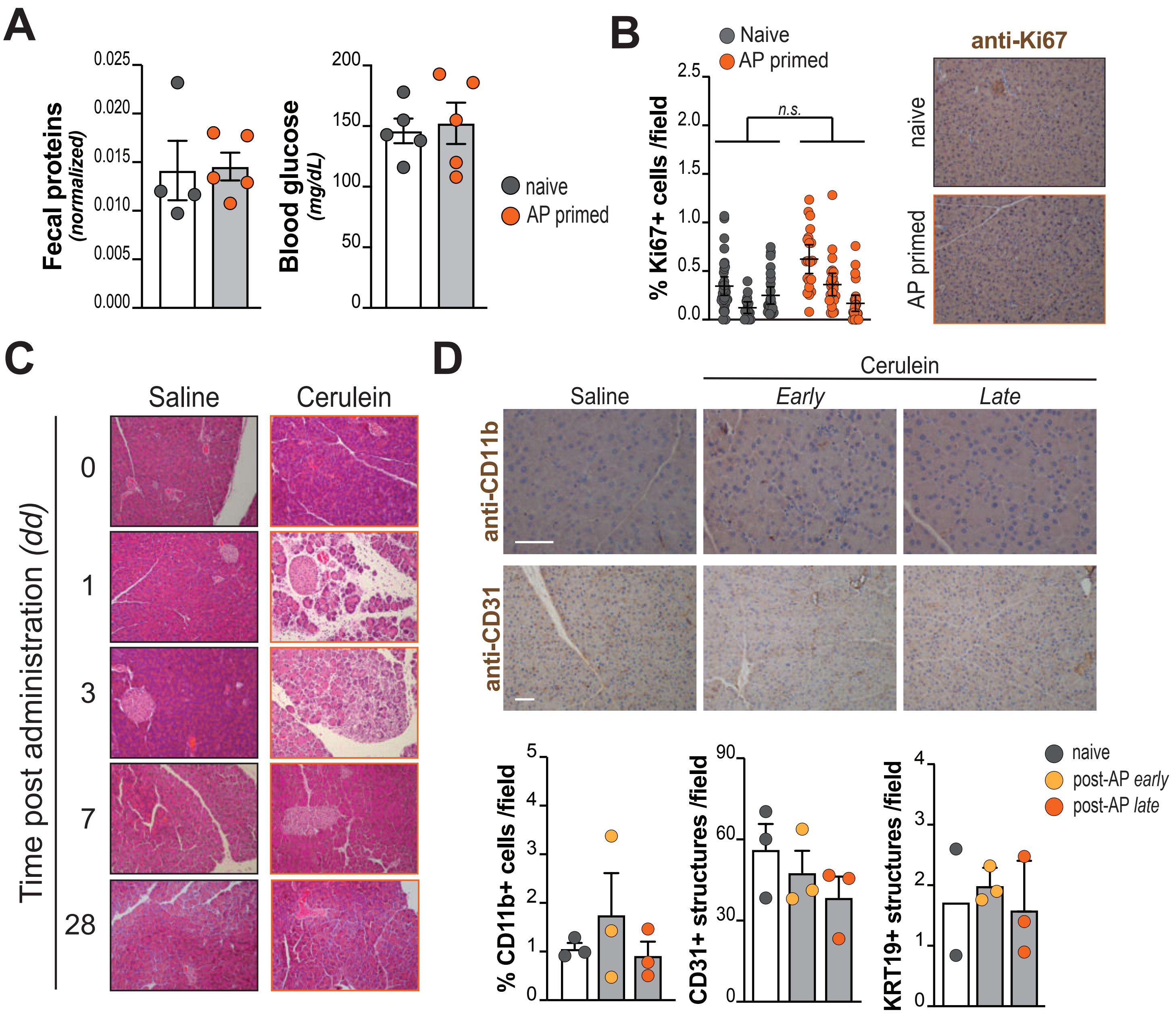

### Supplementary Figure 2

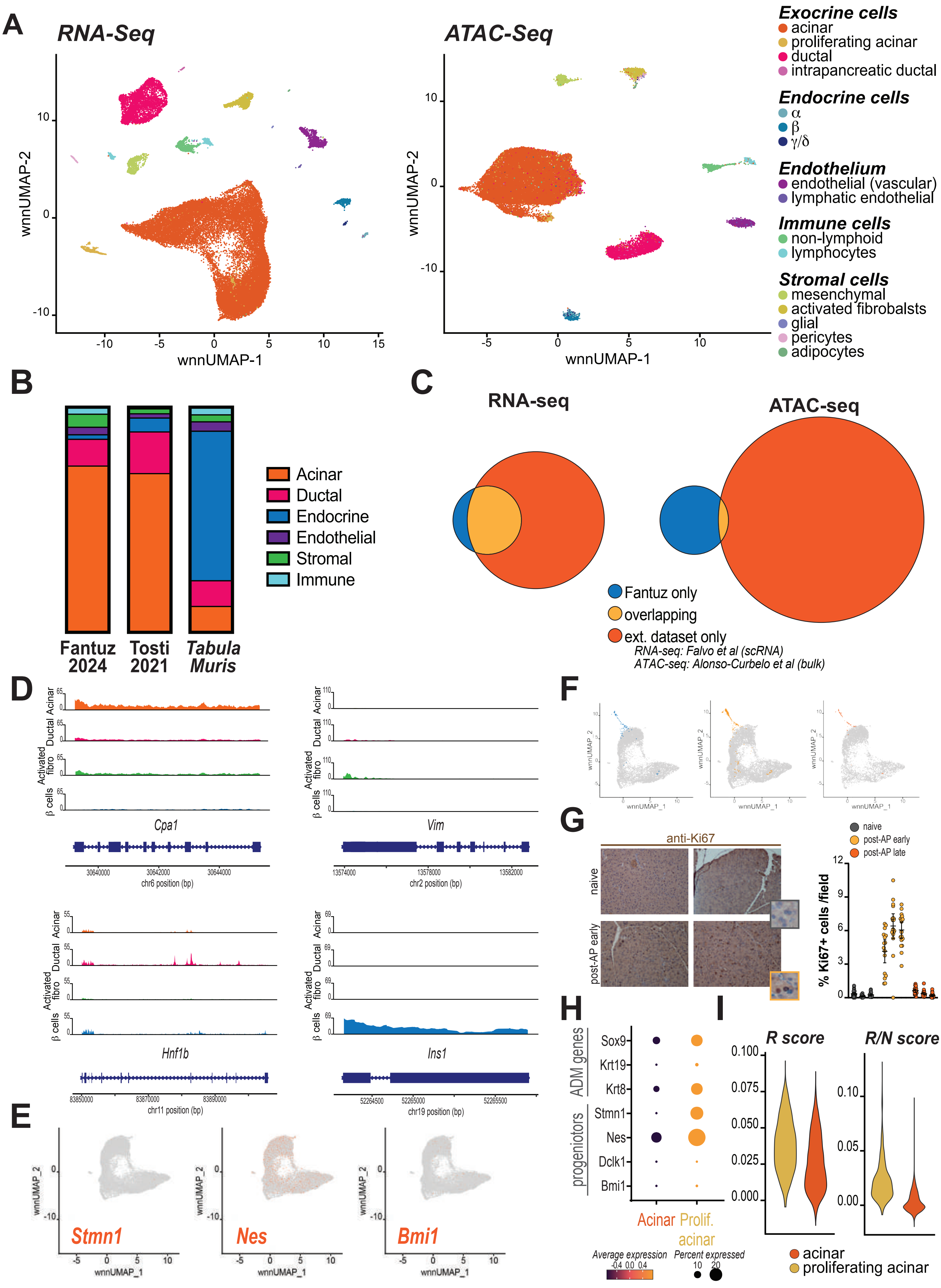

### Supplementary Figure 3

## A

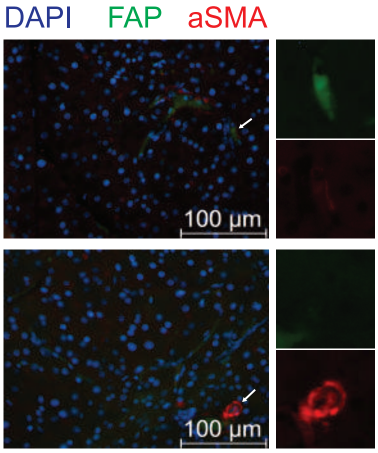

### Supplementary Figure 4

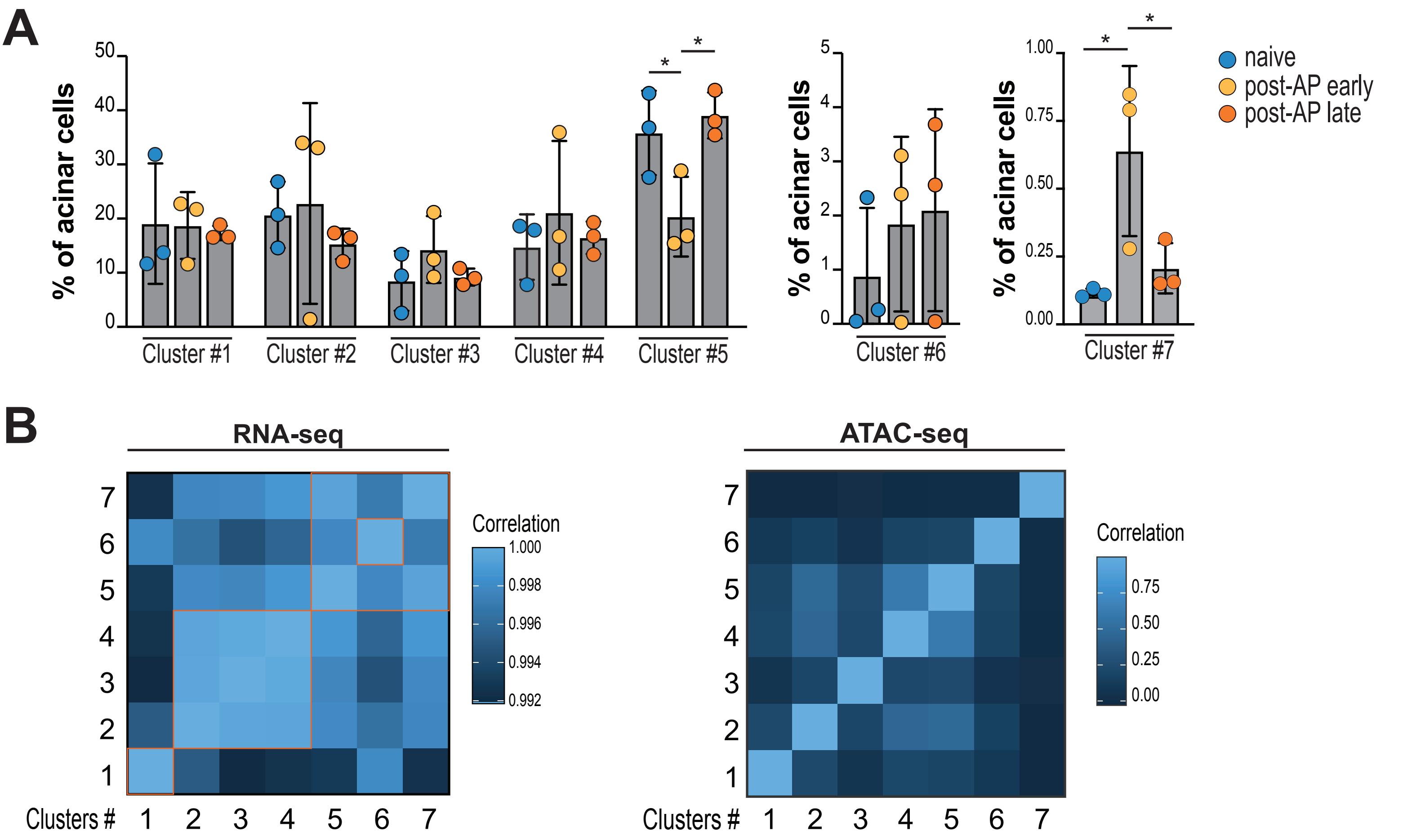

### Supplementary Figure 5

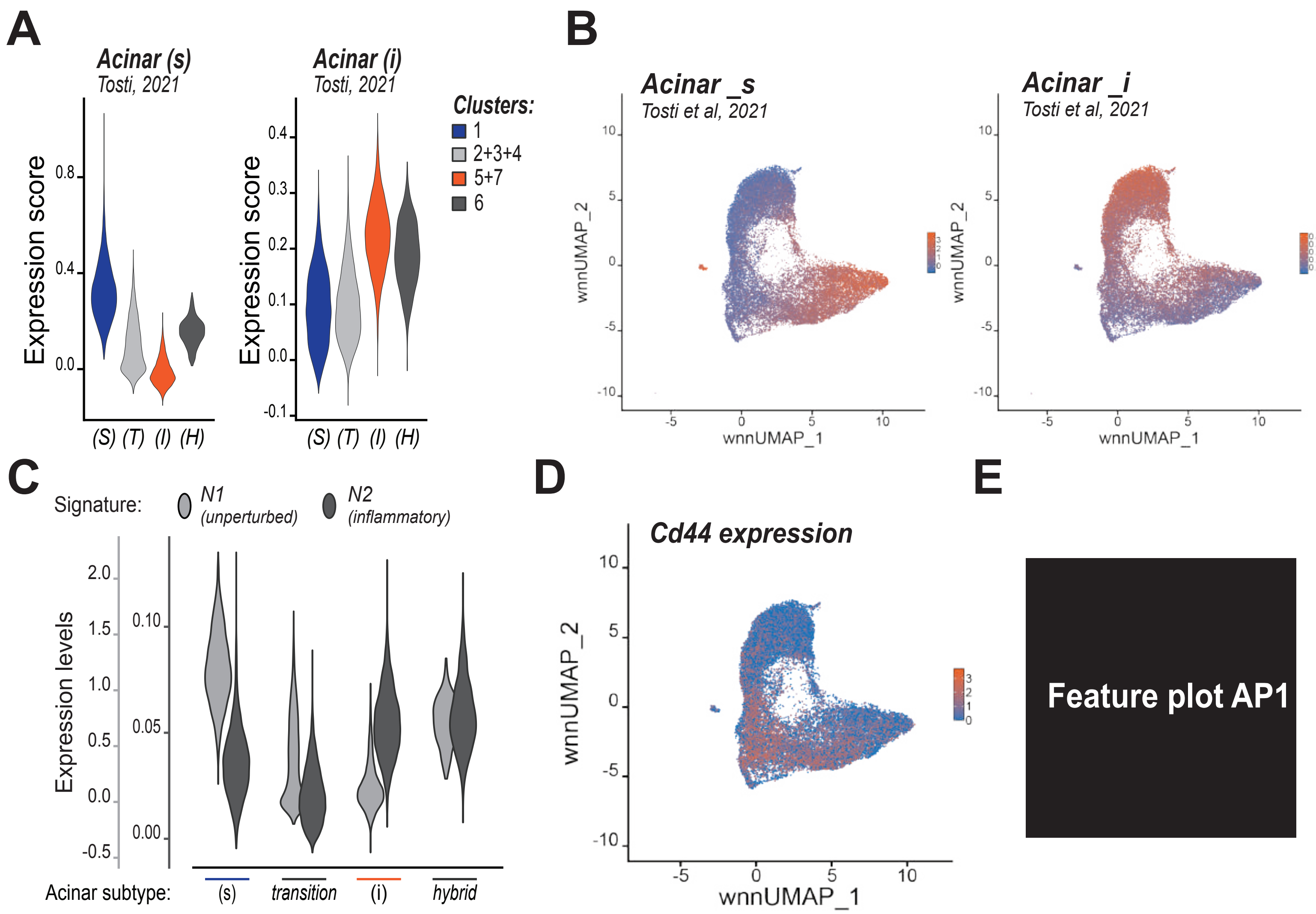

### Supplementary Figure 6

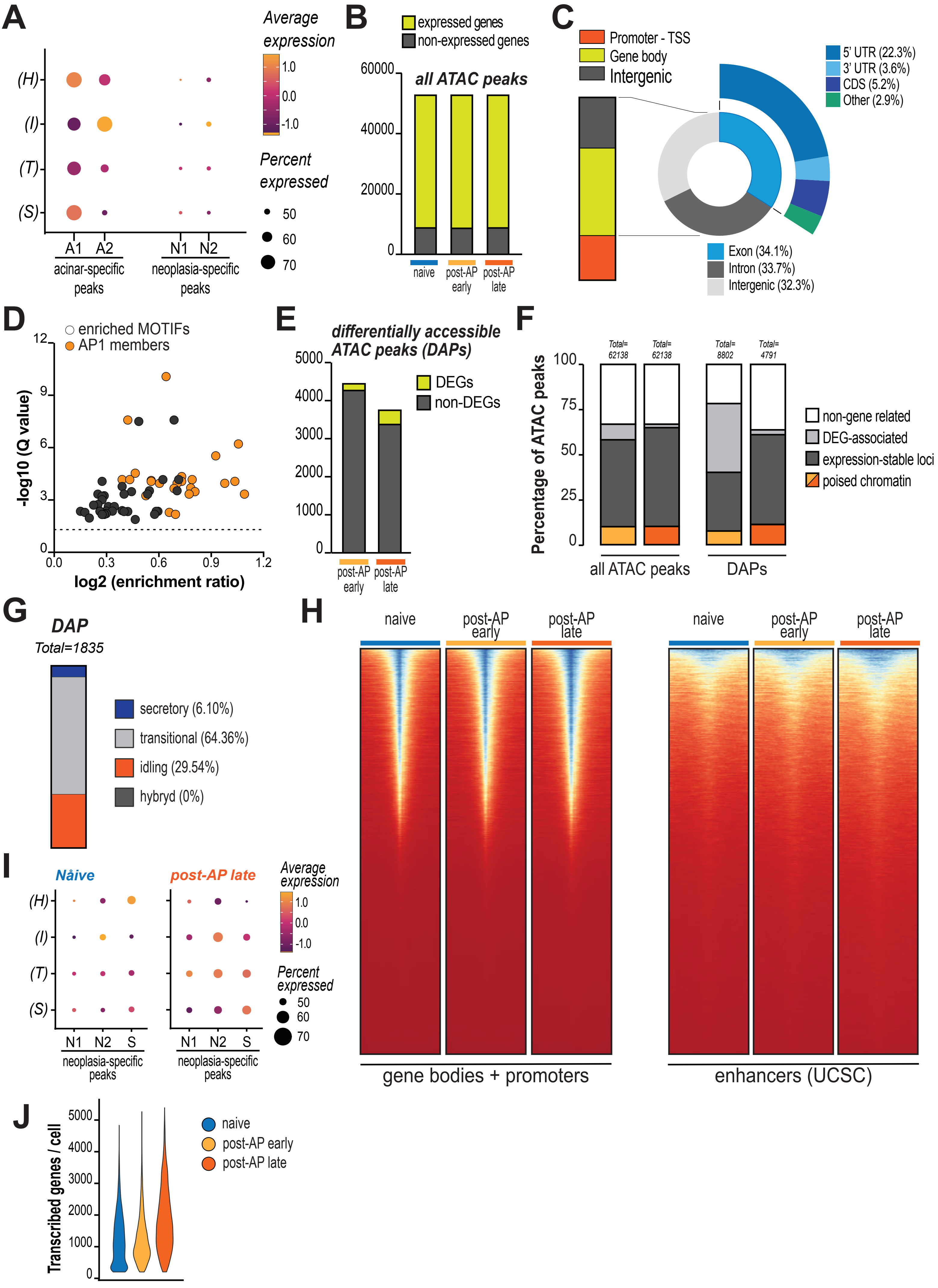

### Supplementary Figure 7

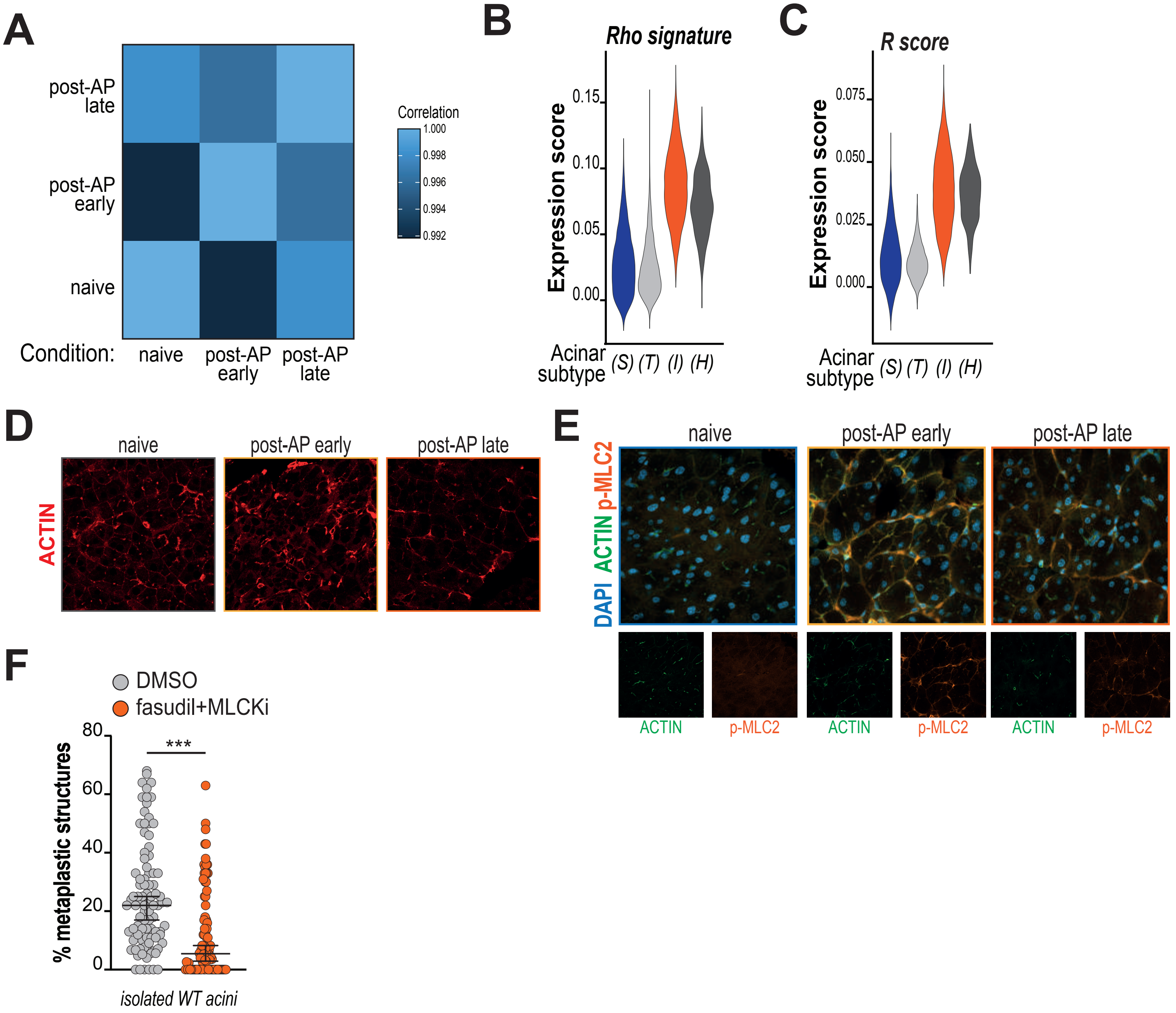

### Supplementary Figure 8

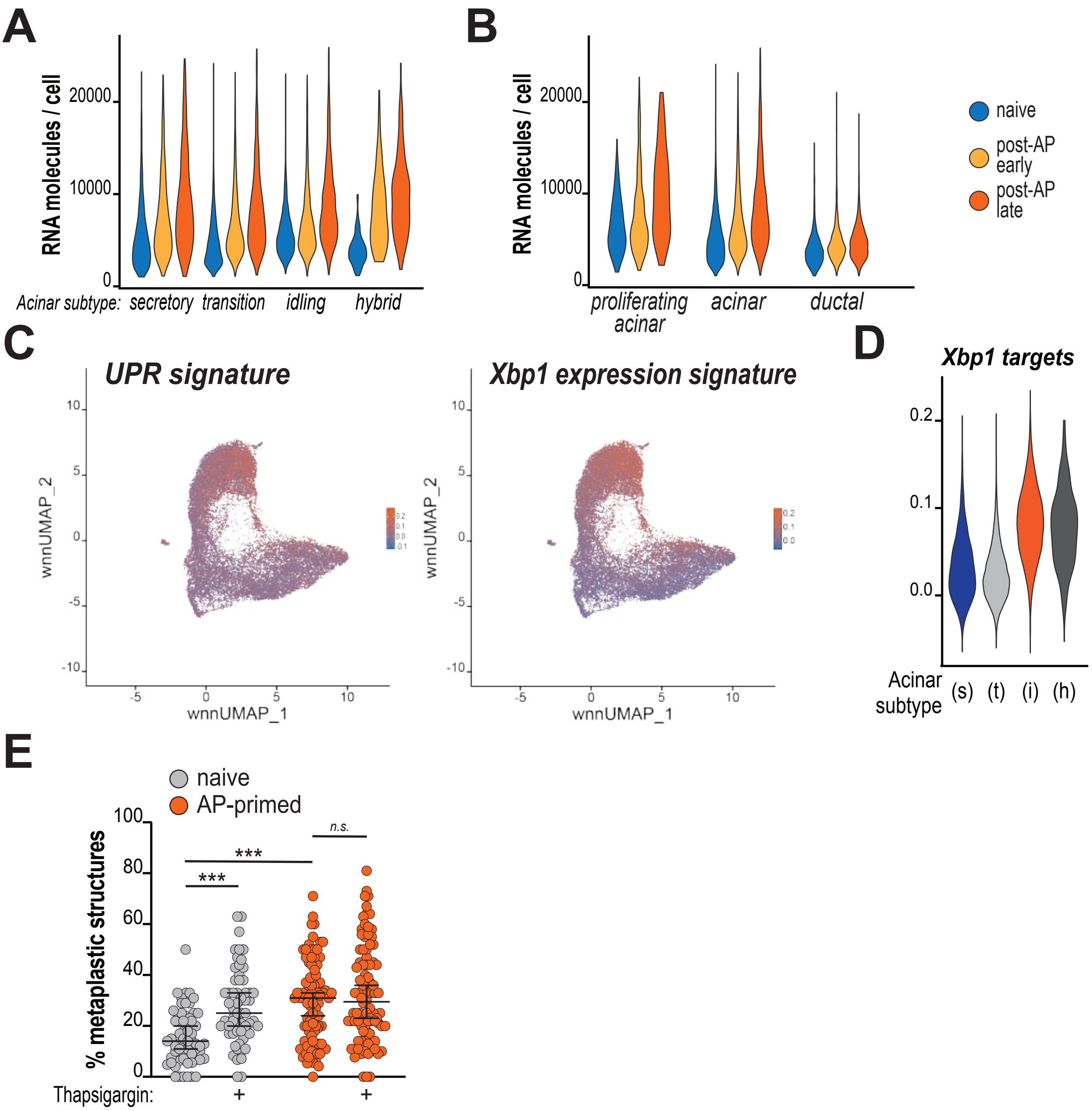
